# SynGAP Splice Variants Display Heterogeneous Spatio-Temporal Expression And Subcellular Distribution In The Developing Mammalian Brain

**DOI:** 10.1101/681148

**Authors:** Gemma Gou, Adriana Roca-Fernandez, Murat Kilinc, Elena Serrano, Rita Reig-Viader, Yoichi Araki, Richard L. Huganir, Cristian de Quintana-Schmidt, Gavin Rumbaugh, Àlex Bayés

## Abstract

The *Syngap1* gene is a major regulator of synapse biology and neural circuit function. Genetic variants linked to epilepsy and intellectual disability disrupt synaptic function and neural excitability. The SynGAP protein has been involved in multiple signaling pathways and can regulate small GTPases with very different functions. Yet, the molecular bases behind this pleiotropy are poorly understood. We hypothesize that different SynGAP isoforms will mediate different sets of functions and that deciphering their spatio-temporal expression and subcellular localization will accelerate our understanding of the multiple functions performed by SynGAP. Using antibodies that detect all isoforms of SynGAP, we found that its subcellular localization changed throughout postnatal development. Consistent with previous reports, SynGAP was enriched in the postsynaptic density in the mature forebrain. However, this was age-dependent and SynGAP was predominantly found in non-synaptic locations in a period of postnatal development highly sensitive to SynGAP levels. Furthermore, we identified different expression patterns in the spatial and temporal axes for different SynGAP isoforms. Particularly noticeable was the delayed expression of SynGAP α1 isoforms, which bind to PSD-95 at the postsynaptic density, in cortex and hippocampus during the first two weeks of postnatal development. The subcellular localization of SynGAP was also isoform-dependent. While, α1 isoforms were highly enriched in the postsynaptic density, other C-terminal isoforms were less enriched or even more abundant in non-synaptic locations, particularly during the postnatal period. Thus, the regulation of expression and subcellular distribution of SynGAP isoforms may contribute to isoform-specific regulation of small GTPases, explaining SynGAP pleiotropy.

## Introduction

*De novo* mutations in the human *SYNGAP1* gene resulting in genetic haploinsufficiency cause an autosomal dominant form of intellectual disability (ID) with high rates of progressively worsening childhood epilepsy (Hamdan *et al*. 2009; Mignot *et al*. 2016; Parker *et al*. 2015; Vlaskamp *et al*. 2019). This debilitating neurodevelopmental disorder is estimated to be responsible for up to 1% of all cases of intellectual disability (Berryer *et al*. 2013). Studies in mouse models of this condition indicate that a *Syngap1* genetic deficit during specific developmental stages causes premature synaptic maturation in excitatory neurons that result in enhanced neuronal excitability (Clement *et al*. 2012; Clement *et al*. 2013; Ozkan *et al*. 2014; Aceti *et al*. 2014). In addition, more recent studies have identified non-developmental functions of the *Syngap1* gene that contribute to memory expression and seizure threshold (Creson *et al*. 2019). Together, these findings indicate that *Syngap1* is critical for brain cell function. Thus, in depth study of this gene will provide insight into the molecular and cellular processes that contribute to neurological and psychiatric disorders.

*Syngap1* encodes the synaptic Ras/Rap GTPase-activating protein (SynGAP), which was first described as one of the most abundant components of the postsynaptic density (Chen *et al*. 1998; Kim *et al*. 1998). Indeed, this protein regulates the structure and function of excitatory synapses in the mammalian forebrain (Kilinc *et al*. 2018; Jeyabalan and Clement 2016). SynGAP has a prominent role in the molecular mechanisms governing synaptic plasticity, being involved in the two hallmarks of this process, incorporation of AMPA receptors into the synaptic plasma membrane (Kim *et al*. 2003; Rumbaugh *et al*. 2006) and dendritic spine enlargement (Vazquez *et al*. 2004; Aceti *et al*. 2014). The activity of SynGAP towards small GTPases is considered to be its key functional role, with the other domains and sequence motifs being involved in regulating it. For instance, the C2 domain is key in the GAP activity towards Rap GTPases (Pena *et al*. 2008) and phosphorylation determines substrate specificity, as CaMKII promotes RapGAP activity while CDK5 and PLK2 stimulate RasGAP activity (Walkup *et al*. 2014; Walkup *et al*. 2018). The exact role of sequences such as the pleckstrin homology (PH) domain, the SH3-binding or poly-histidine motifs in the function of SynGAP are not yet understood. *In vitro* studies with purified proteins have shown that SynGAP directly modulates the activity of HRas (Kim *et al*. 1998), Rap1 (Krapivinsky *et al*. 2004), Rap2 (Walkup *et al*. 2014) and Rab5 (Tomoda 2004). Furthermore, *Syngap1*^+/-^ mice present increased levels of GTP-bound Rac1 in forebrain extracts (Carlisle *et al*. 2008), indicating that SynGAP also regulates Rac1, either directly or indirectly. The GAP activity of SynGAP directly or indirectly participates in the regulation of several important signaling pathways for synaptic physiology, such as Ras-MAPK (Komiyama *et al*. 2002), Ras-PI3K (Qin *et al*. 2005), Rap-p38 (Zhu *et al*. 2002; Krapivinsky *et al*. 2004) and Rac1-PAK (Carlisle *et al*. 2008).

It remains unclear how SynGAP can have such a broad impact on neuronal signaling. Alternative splicing of *Syngap1* mRNA, which results in many protein isoforms, is likely one mechanism. In mammals, the *Syngap1* gene encodes different protein isoforms that differ in their N- and C-terminus (Chen *et al*. 1998; Kim *et al*. 1998; Li *et al*. 2001; McMahon *et al*. 2012). The central part of the protein is thus common to all isoforms and accounts for most of it, extending 1091 residues (>80% of the longest protein isoform) in rat and human. This core region presents a truncated PH domain, lacking the first 24 residues, a C2 domain, a GTPase-activating protein (GAP) domain, a large disordered region of around 600 residues and, finally, a truncated coiled-coil domain lacking its final 11 residues, which is involved in SynGAP trimerization (Zeng *et al*. 2016). Five N-terminal (A1, A2, B, C and D) and four C-terminal (α1, α2, β and γ) SynGAP variants have been described. Of the twenty possible combinations of N- and C-termini with the core region, thirteen have been reported either in NCBI, ENSEMBL or the literature (Chen *et al*. 1998; Kim *et al*. 1998; Li *et al*. 2001). In mouse SynGAP isoforms will vary in their molecular weight, ranging between 148.3 kDa (SynGAP/A2-α2, the largest) and 121.4 kDa (SynGAP/C-β, the smallest). Isoforms with A1/2, B and D N-termini present an entire PH domain, while isoforms containing the C N-terminal do not include its first 24 residues. At the other end of the protein, isoforms with C-terminal variants α1, α2 and γ present an entire coiled-coil domain, while those with the β variant lack its last 11 residues. In support of the idea that *Syngap1* alternative splicing alters protein function, the distinct C-terminal spliced sequences have been shown to cause opposing effects on synaptic strength, with α1 driving synaptic depression (Rumbaugh *et al*. 2006; McMahon *et al*. 2012) and α2 driving synaptic potentiation (McMahon *et al*. 2012).

Thus, the multitude of available N- and C-termini likely bestows distinctive functional properties to SynGAP isoforms. However, the expression pattern and subcellular localization of distinct SynGAP isoforms remain largely unexplored, particularly during early postnatal development, when SynGAP is known to have a strong impact on synaptic (Clement *et al*. 2012; Clement *et al*. 2013) and dendritic (Aceti *et al*. 2014; Michaelson *et al*. 2018) maturation. Here, we present a systematic study of the expression of SynGAP isoforms in five different brain regions and four postnatal developmental stages, identifying specific expression patterns for all isoforms, both between brain regions and throughout development. Furthermore, we investigate the differential subcellular localization of SynGAP isoforms and describe how this varies during cortical development. Together, our data illustrates the complexity of SynGAP roles within brain cells, and the key role that C-term variants are likely to play in SynGAP biology. We also find that SynGAP C-termini are important for its subcellular localization and that SynGAP, generally regarded as almost exclusively found at the synapse, is very abundant in the cytosol, specially early in postnatal development, when the brain is most sensitive to *Syngap1* haploinsufficiency (Clement *et al*. 2012; Aceti *et al*. 2014; Ozkan *et al*. 2014).

## Materials and Methods

### Ethics statement and procedures on human cortical samples

All surgical procedures were approved by the Ethics Committee on Clinical Research from the Hospital de la Santa Creu i Sant Pau (approval reference number 16/041). All samples collected originated from neuro-oncological surgery. Adult healthy cortical samples, as determined by pre-surgery nuclear magnetic resonance, were collected in those cases that a corticectomy had to be performed to access subcortical pathological tissue. All patients were informed and signed an informed consent. Resected tissue was rapidly wrapped in aluminum foil and snap-frozen in liquid nitrogen. Samples were stored in −80°C.

### Ethics statement on animal research and animal handling

All procedures were done with C56BL/6J mice (Jackson laboratories, Main, USA; RRID: MGI:5656552) and in accordance with national and European legislation (Decret 214/1997 and RD 53/2013). These were approved by the Ethics Committee on Animal Research from the Institut de Recerca de I’Hospital de la Santa Creu i Sant Pau (IR-HSCP) and the Departament de Territori i Sostenibilitat from the Generalitat de Catalunya (approval reference num. 9655). Maintenance, treatment and experimental procedures with mice were conducted at the Animal Facility of the IR-HSCP. Mice were housed at a 12h light/dark cycle with fresh water and food *ad libitum*. Special chow (T.2019.12, Envigo) was administered to pregnant mothers and litter until weaning (postnatal day [PND] 21), whereas adult mice were fed with regular chow (T.2014.12, Envigo, Europe). Number of animals used per age: PND4: 28, PND7: 6, PND11: 28, PND14: 6, PND21: 10, PND56: 10. For ages between PND0 and 21, female and male mice were used at equal ratios. PND56 mice were males. Mice culling between PND0 and 4 was performed by head dissection or by cervical dislocation from that age onwards.

### Study time-line

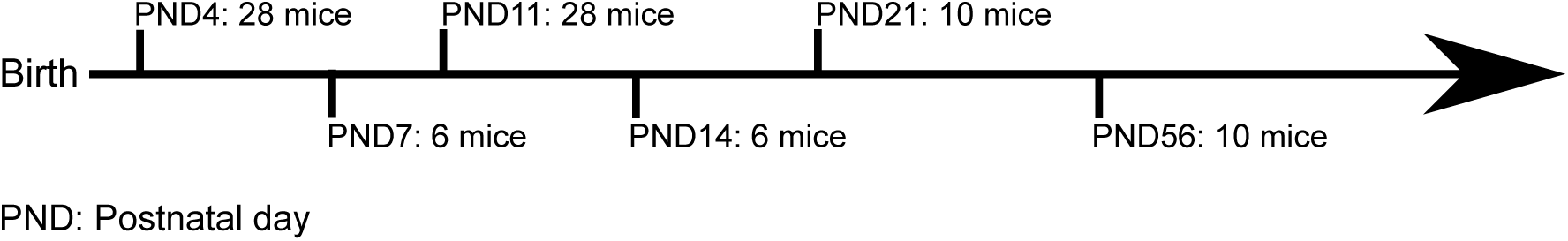

### Mouse brain dissection

Mouse heads were soaked with chilled 1x PBS (0.144 M NaCl; 2.683 KCl mM; 10.144 mM Na2HPO4; 0.735 mM KH2PO4) and dissected using scalpel blades while placed onto a glass petri dish with a filter paper (Merck-Millipore; Darmstadt, Germany). The skull and meninges were removed from brain using Iris scissors and tissue forceps 1:2 (Thermo Scientific). For brain dissection of PND0-7 animals a magnifying loupe (Olympus KC 1500 Ledplus; Olympus, UK) was used. Brain areas were dissected as previously described (Spijker 2011). Tissue weight was recorded before snap-freezing in liquid nitrogen and stored in a −80°C freezer.

### Anti-SynGAP-β antibody generation

SYNGAP beta antibody was raised against SynGAP aa.1273-1285. The antigen peptide with N-terminus Cysteine (NH2-CGGGGAAPGPPRHG-COOH) was coupled with keyhole limpet hemocyanin (Thermo fisher 77600). The antigen was injected into rabbit and antisera were collected after primary and several booster injections. Antisera were further purified with affinity column containing sulfo-link coupling resin (Thermo fisher 20401) coupled with same antigen peptide.

### Total protein extraction, subcellular fractionation and protein quantification

For extraction of total proteins, samples were mixed with chilled buffer (50 mM Tris-HCl pH 9, 1% DOC, 50 mM NaF [Merck-Millipore], 20 mM ZnCl2, 1 mM sodium orthovanadate, 1:2,500 phenyl methane sulfonyl fluoride, 2 µg/mL aprotinin [Merck-Millipore] and 2 µg/mL leupeptin [all from Sigma-Aldrich unless indicated]) at a 1:17.5 tissue:extraction buffer ratio (g/ml). Brain tissue was homogenized by 30 strokes in 1-mL or 7-mL borosilicate Dounce homogenizers (glass-Teflon tissue grinder; Wheaton, Millville, NJ) depending on the volume of buffer required. Then, it was incubated on ice for 1 h and centrifuged at 21,000×g for 30 min at 4 °C in 1.5 mL centrifuge tubes. The resulting pellet was re-homogenized twice following the same procedure and resulting supernatants were pooled. In the last re-homogenization cycle half of the initial w/v ratio was used. Prior to protein quantification, 1% SDS was added to all samples.

Subcellular fractions were prepared following previously described procedures (Carlin *et al*. 1980; Bayés *et al*. 2012). All centrifugation steps were done at 4°C and samples were always kept in ice. Briefly, tissue was homogenized using 7-mL glass-Teflon tissue grinders (borosilicate Dounce homogenizer; Wheaton). A 1:9 ratio was used and ∼40 strokes were applied. Next, a 10 min centrifugation (Epp 5417R, Eppendorf) at 1,400xg was conducted. The resulting supernatant was conserved and the pellet was subjected to two re-homogenizations in the same conditions. The three pooled supernatants were centrifuged at 700xg for 10 min, this sample corresponds with the S1 fraction. This was centrifuged 30 minutes at 21,000xg. The resulting soluble fraction was considered the cytosolic fraction, whereas the pellet obtained contained all membranes. This was resuspended with sucrose 0.32M and 50 mM Tris pH 7.4. A sucrose gradient was prepared with 1 mL of (top to bottom): sample, 0.85 M sucrose and Tris 50 mM pH 7.4; 1 M sucrose and Tris 50 mM pH 7.4, and 1.2 M sucrose and Tris 50 mM pH 7.4, and centrifuged with a SW60 Ti rotor (Beckman Coulter) at 82,500xg for 2h. The interphase between sucrose 1 and 1.2 M was recovered to obtain the synaptosome fraction. The rest of the gradient was centrifuged at 50,000xg 30 min in a fixed rotor and the resulting pellet, containing the non-synaptic membrane (NSM) fraction, was resuspended with 1% SDS and 50 mM Tris pH 7,4. The synaptosome fraction was diluted to reach a final concentration of 10% sucrose with Tris 50 mM pH 7.4 and centrifuged in a Epp 5117R centrifuge (Eppendorff) at 21,000xg during 30 min using 1.5 mL tubes. The resulting pellet was resuspended in Tris 50 mM pH 7.4, 1% Triton X-100 and maintained in ice for 10 min. Finally, samples were centrifuged at 21,000xg during 30 min. As a result, Triton X-100 soluble fraction, referred as synaptic non-PSD (SNP), and the Triton X-100 insoluble enriched in postsynaptic density (PSD), were obtained. Fraction protein yield was defined as the ratio of total protein amount (µg) by tissue weight (mg).

Protein concentration was determined using a micro-BCA protein assay kit (Thermo-Fisher Scientific, Waltham, MA, USA). Prior to IB protein concentrations were corrected by silver stain (Silver Stain Plus™ kit, Bio-Rad).

### Protein dialyzation for detergent exchange

Dialysis was used to exchange DOC with Triton X-100 from total protein extracts prior to immunoprecipitation (IP). Membranes for dialysis (Visking Corporation, Chicago, US) were activated according to manufacturer’s instructions.. Samples were dialyzed ON against the dialysis buffer (50 mM Tris HCl pH7.4 and 1% Triton X-100) at a v/v ratio of 1:1,000 in constant agitation at 4°C. After dialysis, Triton X-100 concentration was adjusted to 1% if required. Finally, samples were sonicated with an ultrasonic bath Sonicator (Thermo Fisher Scientific) at 5% of its maximum intensity during 45 sec with 1.45 sec on/off cycles.

### Immunoprecipitation

IPs were performed on total protein extracts from cortical samples at different ages. All IPs were performed at a protein concentration of 8 mg/mL. All steps were performed at 4°C in an orbital agitator (Stuart). The following amounts of protein were used for each IP: 9 mg for PND0/1 (n of mice = 24), 16 mg for PND11 (n of mice = 15), 8 mg for PND21 (n of mice = 3) and 7 mg for PND56 (n of mice = 3) samples. IPs were performed as described by the kit manufacturer (Pierce® Direct IP Kit, Thermo Fisher Scientific). A sepharose resin (Sigma P3391-250MG) was washed four times with conditioning buffer (50mM Tris pH 7.4). Next a sample pre-clearing step was performed by mixing it with washed resin for 2h at 4°C. Pre-cleared sample was mixed with an anti-SynGAP antibody recognizing an epitope common to all its isoforms (#5540S; Cell Signaling) at a 1:15 (v:v) ratio over-night. Each 200 µL of pre-cleared sample were incubated with 7.5 µL of A sepharose resin during 3h. A 100xg centrifugation step in a column was performed to recover the resin. Resin was washed three times with dialysis buffer and once with conditioning buffer. Bound protein was eluted with 15 µL of the acidic elution buffer from the kit during 10 min.

### Protein electrophoresis

Protein samples for electrophoresis were prepared with Laemmli loading sample buffer (50 mM Tris-HCl, pH 6.8; 2% SDS [Merck-Millipore]; 1% β-mercaptoethanol and 0.04% bromophenol blue) and 10% glycerol (Sigma-Aldrich) and heated at 95°C for 5 min. TGX Stain-Free™ gels (SF gels, Bio-Rad) were prepared and activated according to manufacturer’s instructions. All blue or kaleidoscope precision plus protein standards (Bio-Rad) were used as well as a vertical MiniProtean system kit (Bio-rad) and 1x running buffer (0.025 M TRIS pH 8.4; 0.187 M glycine and 0.1% SDS; all from Sigma-Aldrich). Electrophoretic conditions were 25 mAmp per each 0.75 mm wide gel or 50 mAmp per each 1.5 mm wide gel.

Proteins resolved in SDS-PAGE gels were stained over-night at room temperature with *Coommassie* solution (Bio-Rad). Gels were washed with 2.5 % acetic acid (Sigma-Aldrich) and 20% methanol during 10 min in a rocking platform shaker (Stuart) and later with subsequent washes of 20% methanol, until protein bands were clearly visible. Gel images were acquired with ChemiDoc XRS+ (Bio-Rad) and quantified with Image Studio Lite ver. 3.1 (LI-COR Biosciences).

### Immunoblot

Protein transference was conducted using the MiniProtean kit (Bio-Rad), and 1x chilled transference buffer (20% methanol [Panreac]; 39 mM Glycine; 48 mM TRIS; 0.04% SDS [all from Sigma-Aldrich]). Proteins were transferred into methanol pre-activated polyvinylidene fluoride (PVDF) membranes (Immobilon-P, Merck-Millipore). After transference, PVDF membranes were blocked with 5 mL Odissey blocking solution (LI-COR, Bad Homburg, Germany) prepared with 1x TBS [50mM Tris·HCl pH7.4; 1,5 M NaCl) and 0.1% sodic azide [all from Sigma-Aldrich]) and incubated in a roller mixer (Stuart) with primary antibody solution ON at 4°C. Commercial primary antibodies used were: tSynGAP (NBP2-27541, Novus Biologicals, [RRID: AB_2810282] and Thermo PA1-046, [RRID: AB_2287112], only in Supplementary Figure 2), SynGAP-α1 (06-900, EMD Millipore, [RRID: AB 1163503]), SynGAP-α2 (04-1071 [EPR2883Y], Merck-Millipore, [RRID: AB_1977520]), PSD-95 (3450, Cell Signaling, [RRID: AB_2292883]) and CaMKII-α (05-532, Merck-Millipore, [AB_309787]). Membranes were washed four times with 1x T-TBS for 5 min before incubation for 1 h at RT protected from light with 5 mL of the following secondary antibodies prepared with T-TBS (50mM Tris·HCl pH7.4; 1,5 M NaCl, 0,1% Tween20; all from Sigma-Aldrich): anti-rabbit (926-32313; IRDye 800CW), anti-mouse (926-68073; IRDye 680CW) and anti-goat (926-32214; IRDye 800CW). Membranes were re-blotted without prior stripping by an ON incubation at 4°C or 2h at RT, depending on the antibody. Images were acquired with an Odissey Scanner (LI-COR Biosciences) and protein bands were analyzed with Image Studio Lite ver. 3.1 software (LI-COR Biosciences). Membranes transferred from TGX Stain-Free™ gels were imaged and quantified for posterior normalization steps prior to blocking with a ChemiDoc XRS+ (Bio-Rad) using the Image Lab software (Bio-Rad).

### Normalization of immunoblot data

In spatio-temporal protein expression studies band intensity units (IU) were first corrected for immunoblot technical variability using the value of total protein transferred to PVDF membranes obtained from the TGX Stain-Free^TM^ quantification (IU/protein intensity). Corrected IU were then normalized using the average IU of all bands in a blot. This normalization removed the technical variability between blots allowing to accumulate data from immunoblot replicates.

In subcellular localization studies we first corrected band IU (e.g. tSynGAP in PSD) per amount of total protein used for immunoblotting (e.g. tSynGAP IU in PSD/µg PSD protein). These values were next multiplied by protein yield (with units: µg protein/mg tissue) of their corresponding subcellular fraction, which retrieved a value of specific protein abundance per fraction (e.g. tSynGAP IU in PSD/mg tissue). Finally, these values were normalized by the abundance in the starting homogenate (S1 fraction; e.g. tSynGAP IU in PSD / tSynGAP IU in S1). This normalization step allowed accumulating data from immunoblot replicas.

### Sample preparation and mass spectrometry-based proteomics

Proteins were separated by SDS-PAGE and were stained with *Coomassie* (Bio-Rad). Bands between ∼120-200 kDa were excised from acrylamide gels in a transilluminator (22V, Cultex). Excised gel bands were subjected to an in-gel digestion protocol being first reduced with 10mM dithiothreitol (Sigma-Aldrich) and alkylated with 55mM iodoacetamide (Sigma-Aldrich), and later digested with trypsin (Promega Biotech Ibérica, Madrid, Spain). Tryptic peptides were eluted from acrylamide and around 80% of each trypsin-digested sample was injected in a linear trap quadrupole (LTQ) Orbitrap VelosPro with a short chromatographic method (40min gradient) in a 25 cm 1.9 um column. To avoid carry over, BSA runs were added between samples. BSA controls were included both in the digestion and LC-MS/MS analyses for quality control. This experiment was done twice. The data was searched using an internal version of the search algorithm Mascot (Matrix Science) against a SynGAP (May 2014) homemade database. The Mascot database server search was done with Protein Discoverer ver. 1.4.1.14 (DBVer.:79) using the following search parameters: mass precision of 2ppm; precursor mass range of 250 Da to 5,000 Da; Trypsin with a maximum of 3 miss-cleavages; the peptide cut off score was set at 10 and peptide without protein cut off at 5. Peptides were filtered based on IonScore>20. The precursor mass tolerance (MS) was set at 7 ppm and fragment mass tolerance (MS/MS) at 0.5 Da with two variable modifications: oxidation (M) and acetylation (protein N-term), and one fixed modification (C): carbamidomethyl. False discovery rates (FDR) determined by reverse database searches and empirical analyses of the distributions of mass deviation and Mascot Ion Scores were used to establish score and mass accuracy filters. Application of these filters to this dataset was below 1% FDR as assessed by reverse database searching.

### Data statistical analyses

Statistical tests used are indicated in figure legends, together with the exact number of biological and technical replicates. GraphPAD Prism version 6.0 (GraphPad, San Diego, CA) was used to conduct statistical analyses. When required data was assessed for normal distribution by descriptive statistic measures (mean and median) and applying the Shapiro-Wilk and Kolmogorov-Smirnov tests. All statistical analyses were conducted with a significance level of α = 0.05 (p ≤ 0.05). No blinding was performed. No statistical method was used to determine sample size, which was determined based on the previous experience of the group with the goal to minimize the number of animals required.

## Results

### Total SynGAP protein expression is different between brain regions and changes throughout postnatal development

Using an antibody that recognizes a sequence common to all SynGAP isoforms we have analyzed by immunoblot the abundance of all of them together, what we have called total SynGAP (tSynGAP). We have investigated tSynGAP expression in five mouse brain regions (cortex, hippocampus, striatum, olfactory bulb and cerebellum) at 4, 11, 21 and 56 postnatal days (PND) of life. Depending on the tissue, tSynGAP presents three patterns of developmental expression (Fig.1A). In cortex and hippocampus, tSynGAP increases sharply, reaching its maximum at PND21, and remaining at this level until PND56. Between PND4 and PND21, tSynGAP levels increase over six times in both tissues. Striatum presents a different pattern. tSynGAP expression is maintained constant between PND11 and PND21, and its maximum level is not reached until PND56. Yet tSynGAP levels also increase notably, also around six times, between PND4 and 56. Finally, both the olfactory bulb (OB) and the cerebellum present a very modest, albeit significant, increase of tSynGAP levels. Between PNDs 4 and 56, tSynGAP increases 1.7 times in OB and 1.2 in cerebellum.

**Figure 1.**
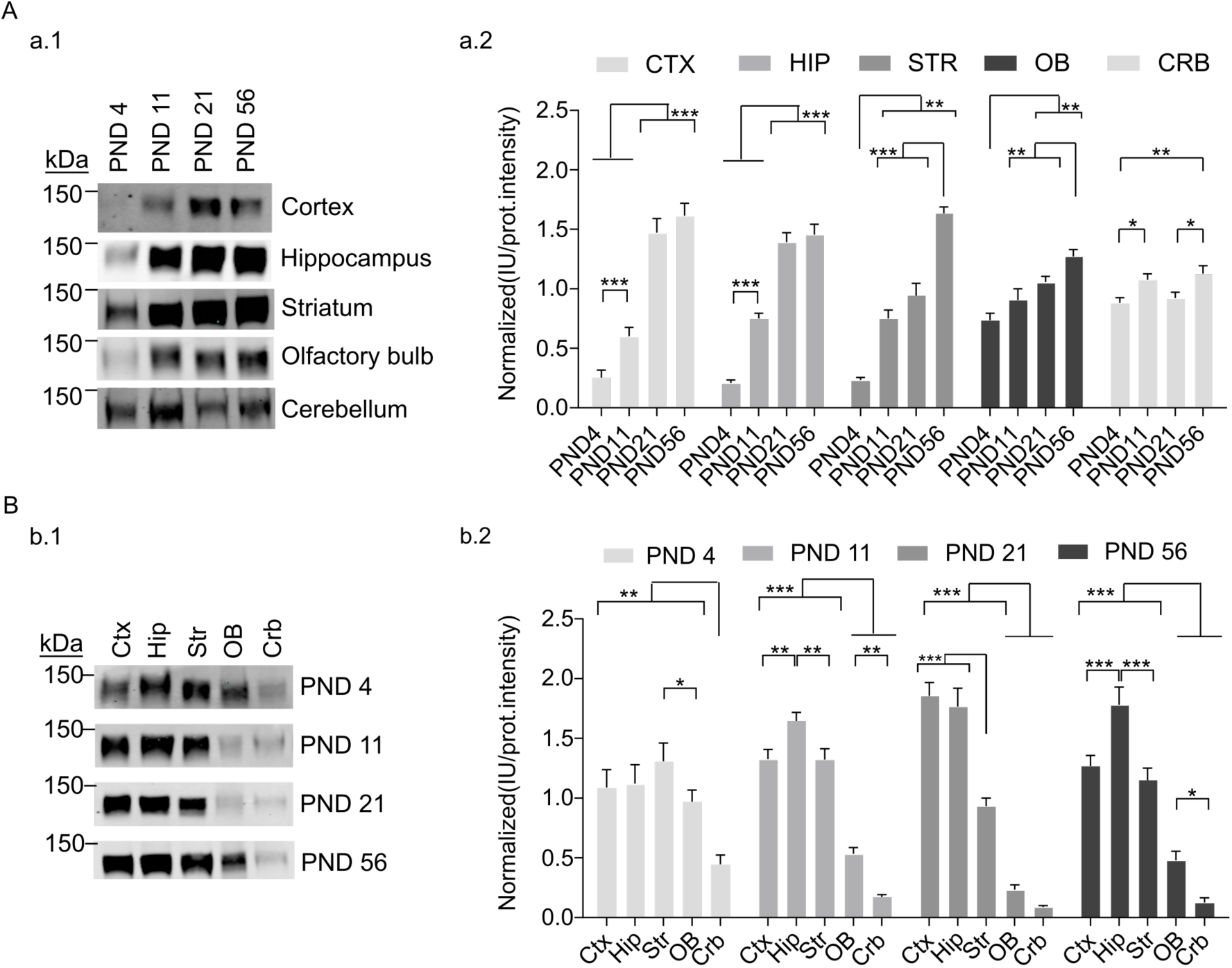
Abundance of tSynGAP in five different brain regions and four postnatal stages. **A.** Developmental changes in tSynGAP abundance in five different brain regions (cortex, hippocampus, striatum, olfactory bulb and cerebellum). Ages investigated were postnatal day (PND) 4, 11, 21 and 56. **a.1.** Representative immunoblots showing tSynGAP abundance in each of the five tissues. **a.2.** Bar plots depict the mean of normalized protein abundance data derived from immunoblot intensity (N: cortex 14-20, hippocampus 12-16, striatum 6-7, olfactory bulb 11-16 and cerebellum 23-24). The standard error of the mean (SEM) is also shown. Mean differences were analyzed by one-way ANOVA followed by Tukey’s post-hoc test, *** p < 0.001, ** p < 0.01 and * p < 0.05. **B.** Brain region changes in tSynGAP abundance in four life stages, including three postnatal development stages (PND4, 11 and 21) and adulthood (PND56). **b.1.** Representative immunoblots showing tSynGAP abundance in each life stage. **b.2.** Bar plots depict the mean of normalized protein abundance data derived from immunoblot intensities (N: cortex 6-15, hippocampus 6-15, striatum 6-15, olfactory bulb 6-15 and cerebellum 6-15). The standard error of the mean (SEM) is also shown. Mean differences were analyzed by one-way ANOVA followed by Tukey’s post-hoc test, *** p < 0.001, ** p < 0.01 and * p < 0.05.

We have also investigated how tSynGAP levels compare between tissues at each of these four developmental stages (Fig. 1B). Early in postnatal development tSynGAP levels are very similar in cortex, hippocampus, striatum and olfactory bulb, while cerebellum already presents the lowest levels. At PND4 there is approximately 2.5 times more tSynGAP in forebrain regions than in cerebellum. This difference becomes larger with age, reaching a maximum difference of 30 times when comparing hippocampal and cerebellar expression at PND21/56 or cortical expression at PND21. As mice develop, the levels of tSynGAP in olfactory bulb also lag behind those of the other forebrain areas (Fig. 1B), being the maximum difference at PND21, when cortex and hippocampus express 7 times more tSynGAP than olfactory bulb. At PND11, cortex and striatum display similar levels of tSynGAP, while hippocampus presents a slight, but significantly higher abundance, being the tissue with the highest tSynGAP levels at this age. At PND21, cortical and hippocampal tSynGAP have similar levels, presenting almost twice as much tSynGAP than striatum. This is in agreement with the sustained tSynGAP levels previously observed in striatum between PND11 and 21 (Fig. 1A). Finally, at PND56, the abundance profile of tSynGAP at cortex, hippocampus and striatum is very similar to that found at PND11. Cortex and striatum have similar abundance, while hippocampus presents significantly more tSynGAP.

### *In silico* identification of novel *Syngap1* splice variants

ENSEMBL, NCBI-Gene and UniProt (as of 02 May 2019), together with the previous literature (Chen *et al*. 1998; Kim *et al*. 1998), report a total of 15, 9 and 7 *Syngap1* transcripts in mouse, rat and human, respectively (Suppl. Table 1). Remarkably, there is still little overlap between these databases. For instance, in mouse, only 2 proteins can be directly related between ENSEMBL and NCBI. Interestingly, the NCBI Gene database identifies unpublished variants in mice (four N-terminal and one C-terminal). We refer to these unreported N-terminals as A3, A4, E and F and to the C-terminal one as α3 (Suppl. Fig. 1). The first two N-term variants are shorter versions of A1/2, E presents a unique N-terminus and F starts at residue 430 inside the core of SynGAP. If the F variant is expressed at the protein level, it would lack the PH, C2 and GAP domains, which could be functionally relevant to neuronal biology. In order to investigate if any of these predicted variants is expressed at the protein level, we immunoprecipitated tSynGAP from mouse cortex at four postnatal stages and performed high-throughput mass spectrometry-based proteomics. However, we could not identify unique peptides for any of these variants. Instead, we identified a unique peptide corresponding to the first residues of the D N-terminus (Suppl. Table 2), which had only been reported at the RNA level (Li *et al*. 2001). Importantly, this peptide presented an acetylated initial methionine, which is a common post-translational modification of the N-terminus. Considering that we identified this acetylated peptide in mouse cortex at four different postnatal stages, it is highly likely that this sequence corresponds to a novel SynGAP N-terminus, expressed at least in mouse cortex.

### SynGAP isoforms present different developmental expression patterns

SynGAP isoforms present four different C-terminal variants that have been identified at the protein level, which are named alpha1 (α1), alpha2 (α2), beta (β) and gamma (γ). Commercial antibodies are available for two (α1 and α2) and we raised a new antibody that recognizes the β sequence. We confirmed that these three antibodies are selective by showing that they do not cross-react (Suppl. Fig. 2). In our experimental conditions, as in previous works (Li *et al*. 2001; Yang *et al*. 2013; Kim *et al*. 1998; McMahon *et al*. 2012), these C-terminal specific antibodies distinguish two major bands (Figs. 2-3). These two bands correspond with at least two different isoforms, which will necessarily present different N-terminus. Notably, we have not found statistically significant abundance differences between the top and bottom bands in any of the experiments performed. This indicates that isoforms with the same C-terminus display equivalent expression patterns along development in the five brain regions investigated. For this reason, we considered both bands together for subsequent analysis.

In cortex (Fig. 2A), α1-containing SynGAP isoforms remain at very low levels until PND11 as compared with their maximum expression. Between PND11 and PND21, α1 expression increases five-fold, to reach over 60% of their adult (PND56) levels. In contrast, isoforms containing α2 and β C-term variants already present around 50% of their maximum abundance at PND11. Interestingly, α1-, α2- and β-containing isoforms vary in their pattern of cortical expression. Namely, α1 isoforms do not reach their maximum until PND56, while α2 and β isoforms peak at PND21. Furthermore, while α2 isoforms maintain their maximum expression level between PND21 and 56, those of β isoforms decrease significantly after PND21, presenting 70% of their maximum expression at PND56.

**Figure 2.**
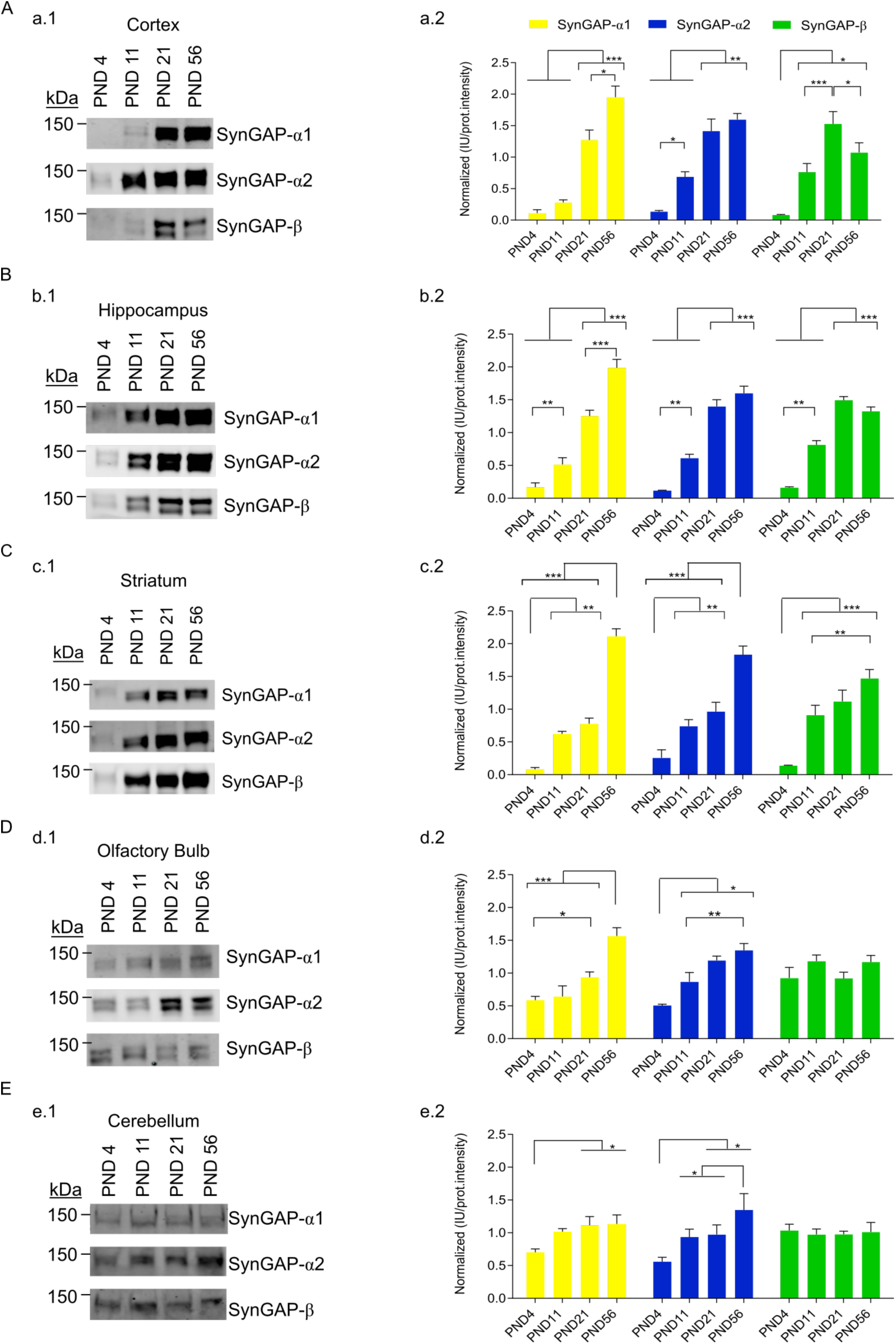
Compared protein abundance of SynGAP isoforms along postnatal development in five different brain regions. **A** to **E** data from cortex, hippocampus, striatum, olfactory bulb and cerebellum, respectively. **a1** to **e1**, representative immunoblots for SynGAP isoforms containing each of the three C-terminal variants (α1, α2 and β). **a2** to **e2**, bar plots for each brain region depicting mean normalized protein abundance data from each isoform derived from immunoblot intensities (N: cortex 4-19, hippocampus 6-12, striatum 3-9, olfactory bulb 6-14 and cerebellum 8-15). The standard error of the mean (SEM) is also shown. Mean differences were analyzed by one-way ANOVA followed by Tukey’s post-hoc test, *** p < 0.001, ** p < 0.01 and * p < 0.05.

The hippocampal expression pattern (Fig. 2B) of the SynGAP isoforms investigated is quite similar to that of cortex. α1 isoforms reach their maximum expression at PND56, while α2 and β isoforms peak at PND21. Here we also observed a decrease of β isoforms between PND21 and 56, although it did not reach statistical significance. Also, the abundance of α1 isoforms does increase between PND4 and 11, as opposed to what we observed in cortex. Still, α1 isoforms expression fold change between PND11 and PND56 is higher (4-fold) than that obserbed between PND4 and 11 (2-fold).

In striatum (Fig. 2C), α1 and α2 isoforms present a biphasic expression pattern that we have not observed in any other tissue. Expression increases from PND4 to PND11 and then again between PND21 and PN56, but during the second and third weeks (PND11-21) the expression of these isoforms remains constant. Striatal levels of β isoforms suggest a similar pattern, as PND56 expression is higher than PND11 and there is no difference between PND11 and PND21. Yet, the difference between PND21 and PND56 does not reach statistical significance. Thus, our data could also be interpreted as that β isoforms reach their maximum level at PND21 and then this is maintained.

In the olfactory bulb and cerebellum (Fig. 2D,E) we observed less developmental variation in the abundance of SynGAP isoforms. Particularly for β isoforms, which do not present any difference in their expression along the postnatal period investigated. In olfactory bulb, both α1 and α2 isoforms present a moderate increase in their expression level, showing a maximum at PND56. Finally, in cerebellum, α1 levels are constant from PND11 onwards, while α2 present a biphasic increase in expression, with a period of latency between PND11 and PND21.

### SynGAP isoforms present different regional expression patterns

We next compared the abundance of SynGAP isoforms between brain regions in three postnatal developmental time points, PND4, 11, 21 and in young adults (PND56). At PND4 (Fig. 3A), the expression of all isoforms investigated presented equivalent levels in cortex, hippocampus and striatum, while cerebellar expression was always the lowest. In the olfactory bulb, α2 and β levels were undistinguishable from those in the other forebrain areas, but α1 abundance was significantly reduced, presenting the same levels found in cerebellum. Actually, α1 isoforms present similarly low expression levels in cerebellum and olfactory bulb in all ages investigated. At PND11, α2 and β-containing isoforms still present higher levels in olfactory bulb when compared with cerebellum, yet this difference disappears at PND21 and 56, where both tissues express equally low levels of all isoforms.

**Figure 3.**
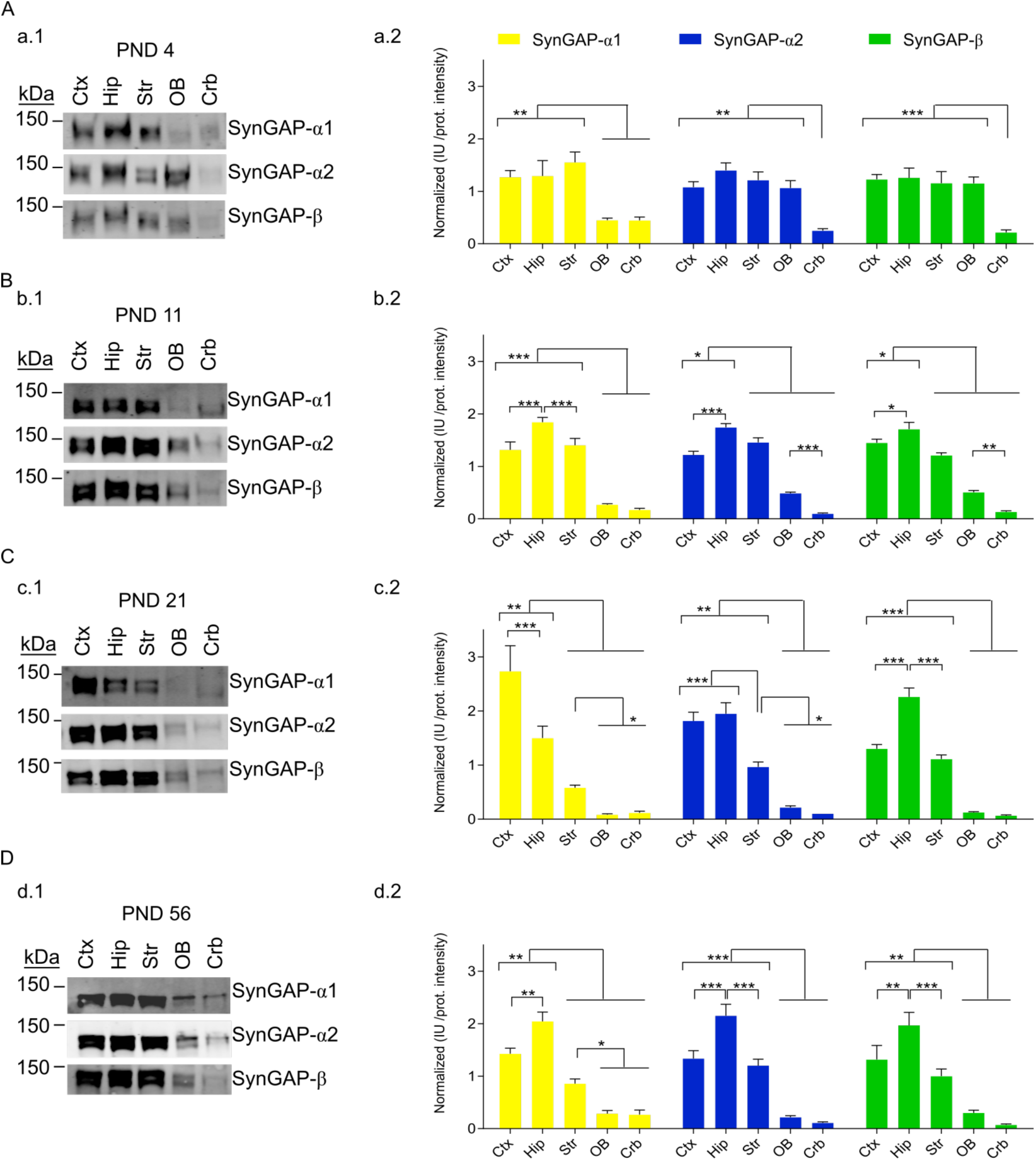
Compared protein abundance of SynGAP isoforms in five different brain areas along postnatal development and in adulthood. **A** to **D** data from postnatal day (PND) 4, 11, 21 and 56, respectively. **a1** to d**1**, representative immunoblots for SynGAP isoforms containing each of the three C-terminal variants (α1, α2 and β). **a2** to d**2**, Bar plots for each life stage depicting normalized protein abundance data from each isoform derived from immunoblot intensities (N: cortex 6, hippocampus 6, striatum 6, olfactory bulb 6 and cerebellum 6). The standard error of the mean (SEM) is also shown. Mean differences were analyzed by one-way ANOVA followed by Tukey’s post-hoc test, *** p < 0.001, ** p < 0.01 and * p < 0.05.

After PND4, hippocampus was the region where isoforms presented the highest levels (Fig. 3B-D). This was already noticeable at PND11, although not all comparisons reached statistical significance, and very clear at PND56. Nevertheless, at PND21 cortical expression of α1 and α2 isoforms becomes more prominent. This phenomenon is particularly noticeable for the cortical expression of α1 isoforms, which present a cortex to hippocampus expression ratio close to 2 at PND21, and around 0.7 at PND11 and PND56. The cortical expression of α2 isoforms also increases, although not so much, as they present the same expression levels found in hippocampus.

### Expression correlation between SynGAP isoforms and their interactors

SynGAP α1 isoforms interact with PSD-95 (Kim *et al*. 1998) and β ones with CaMKIIα (Li *et al*. 2001). Therefore, we investigated the developmental expression profiles of PSD-95 and CaMKIIα (Fig 4B) and compared them to that of α1 and β isoforms, respectively. Interestingly, we found a positive and statistically significant correlation for the expression of PSD-95 and SynGAP α1 isoforms in cortex, hippocampus and olfactory bulb, but not in cerebellum. The correlation for striatum was also high (0.94), although it did not reach statistical significance (p =0.06, Fig. 4C,D). The main difference in striatal expression difference between these two proteins is that PSD-95 expression is not halted between PND11 and 21. Instead, the developmental expression profile of α1 isoforms and CaMKIIα did not significantly correlate in any of the tissues investigated. As expected, the correlation between SynGAP α1 and β isoforms was also low. In contrast, the expression of SynGAP β isoforms correlated with that of CaMKIIα, but only in cortex and hippocampus. We also observed that β isoforms present good expression correlation with PSD-95, although they do not directly bind to it. These data suggest that the expression between β-containing SynGAP isoforms and CaMKIIα might be less specific than the one between α1 isoforms and PSD-95.

**Figure 4.**
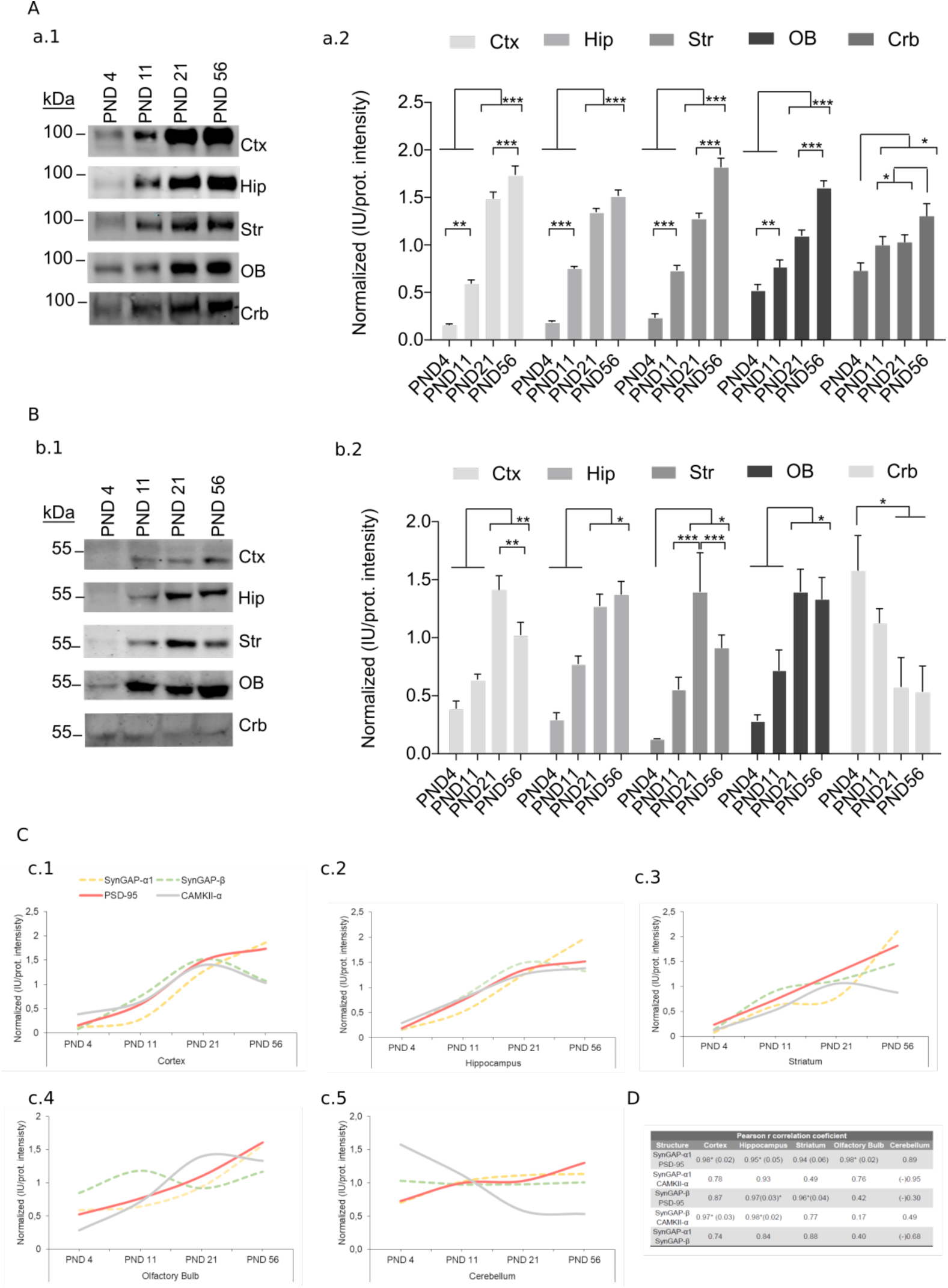
PSD-95 and CaMKIIα protein abundance along postnatal development in five different brain regions. **A.** PSD-95 protein abundance in four life stages (PND4, 11, 21 and 56) and five brain regions; cortex (Ctx), hippocampus (Hip), striatum (Str), olfactory bulb (OB) and cerebellum (Crb). **a.1.** Representatives immunoblots for each of the brain areas investigated. **a.2.** Bar plots represent the mean of normalized immunoblot intensity data (N: cortex 17-20, hippocampus 7-12, striatum 15-19, olfactory bulb 18 and cerebellum 11-13). The standard error of the mean (SEM) is also shown. Mean differences were analyzed by one-way ANOVA followed by Tukey’s post-hoc test, *** p < 0.001, ** p < 0.01 and * p < 0.05. **B.** CaMKIIα protein abundance in four life stages (PND4, 11, 21 and 56) and five brain regions; cortex (Ctx), hippocampus (Hip), striatum (Str), olfactory bulb (OB) and cerebellum (Crb). **b.1.** Representative immunoblots for each of the brain areas investigated. **b.2.** Bar plots represent the mean of normalized immunoblot intensity data (N: cortex 11-20, hippocampus 7-12, striatum 3-11, olfactory bulb 7-11 and cerebellum 3-4). The standard error of the mean (SEM) is also shown. Mean differences were analyzed by one-way ANOVA followed by Tukey’s post-hoc test, *** p < 0.001, ** p < 0.01 and * p < 0.05. **C-G.** Summary graphs representing abundance of SynGAP-α1 and –β isoforms together with that of PSD-95 and CaMKIIα. Each graph presents the data from one brain region as indicated. **H.** Table with correlation values for developmental protein expression of SynGAP-α1 and –β isoforms, PSD-95 and CaMKIIα in graphs C-G. Data passed Shapiro-Wilk and Kolmogorov-Smirnov normality tests and the Pearson correlation coefficient was computed. For statistical significances, paired t-test was applied * p<0.05.

### All SynGAP isoforms investigated are expressed in the young and old human cortex

Immunoblot of total cortical extracts from two human individuals with 19 and 67 years of age was performed to elucidate if SynGAP isoforms are expressed in this human tissue. We found expression of tSynGAP and all C-terminal variants in both samples (Fig. 5). In line with the lower neuronal density found in the human cortex, as compared with the mouse one (Defelipe *et al*. 2002), human immunoblots present a clear reduction per unit of protein of tSynGAP and all its isoforms. Interestingly, human samples presented the same two bands found in mice when investigated by immunoblot. The two human samples analyzed suggest that there is a decrease in the amount of all SynGAP isoforms with age. However, further experiments will be required to demonstrate this initial observation.

**Figure 5.**
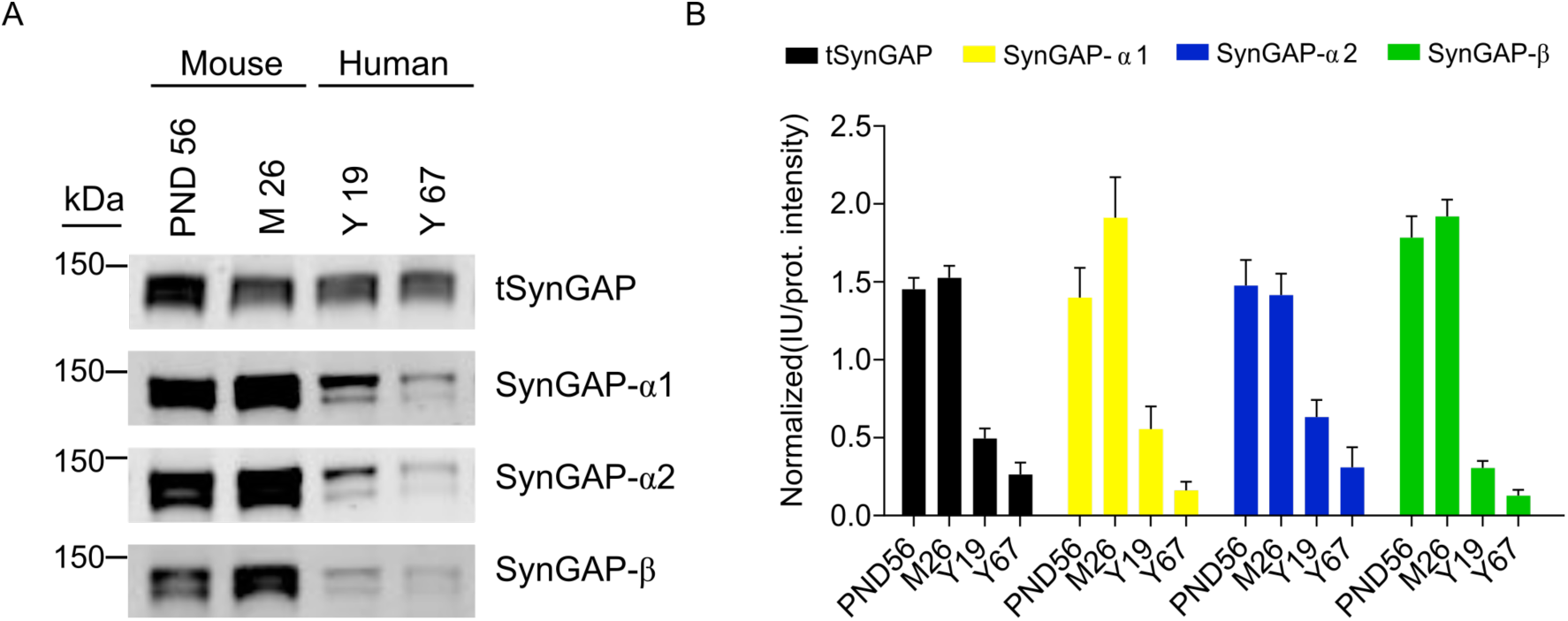
Expression of SynGAP isoforms in human cortex. **A.** Representative immunoblots showing expression of total SynGAP (tSynGAP) as well as α1-, α2- and β-containing SynGAP isoforms in total protein extracts from human cortex. Two samples from human individuals aged 19 and 67 years were investigated. For comparison mouse cortex homogenates from PND56 and 26 months of age animals (M26) were analyzed. **B.** Bar plots represent the mean of normalized immunoblot intensity data (N: mouse PND56 9-26, mouse 26 months 9-23, human 19 years 10-24, human 67 years 9-16). The standard error of the mean (SEM) is also shown. No statistics were performed as we only used 2 human samples.

### Differential subcellular distribution of tSynGAP along postnatal development

Next, we investigated to what extent tSynGAP presented a differential subcellular distribution throughout postnatal development. This study was performed with mouse cortical samples. Four subcellular fractions were prepared: cytosol, non-synaptic membranes, synapse-not-PSD (SNP) and PSD. This was done for three postnatal development stages PND7, 14 and 21 and young adult (PND56). Protein yield (i.e., protein amount / tissue weight) was calculated for all subcellular fractions generated (Suppl. Fig. 3A) and used, together with immunoblot intensity data (Suppl. Fig. 3B), to obtain a measure of protein abundance in each fraction. Interestingly, between PND14 and 21, we observed a decrease in the SNP yield simultaneous to an increase in the PSD yield, likely reflecting the increased maturation of synapses during this week (Suppl. Fig. 3C).

Abundance of tSynGAP in different subcellular fractions was analyzed by immunoblot together with that of PSD-95, a very well established marker of PSDs (Fig. 6A-D). Of the four subcellular fractions investigated, tSynGAP was essentially found in two, cytosol and PSD. Indicating that tSynGAP is not associated to extra-synaptic membranes and that within the synapse, it is essentially found at the PSD. PSD-95 expression always presented a very restricted expression at the PSD. Interestingly, at PND7 tSynGAP was largely found in the cytosol and it is not until PND56 that we observed significantly more tSynGAP in the PSD than in the cytosolic fraction. At PND14 and 21, the abundance of tSynGAP at the PSD was significantly higher than that found in the SNP fraction, yet tSynGAP was still more abundant in the cytosol. At PND21, the cytosol:PSD ratio of tSynGAP is 7:4. Even at PND56, the remaining fraction of tSynGAP at the cytosol is quite large, as this ratio is 5:7. This developmentally regulated subcellular expression pattern of tSynGAP is in stark contrast with that displayed by PSD-95, as this scaffolding protein is predominantly expressed at the PSD in all life stages investigated, presenting almost negligible levels in the cytosolic fraction.

**Figure 6.**
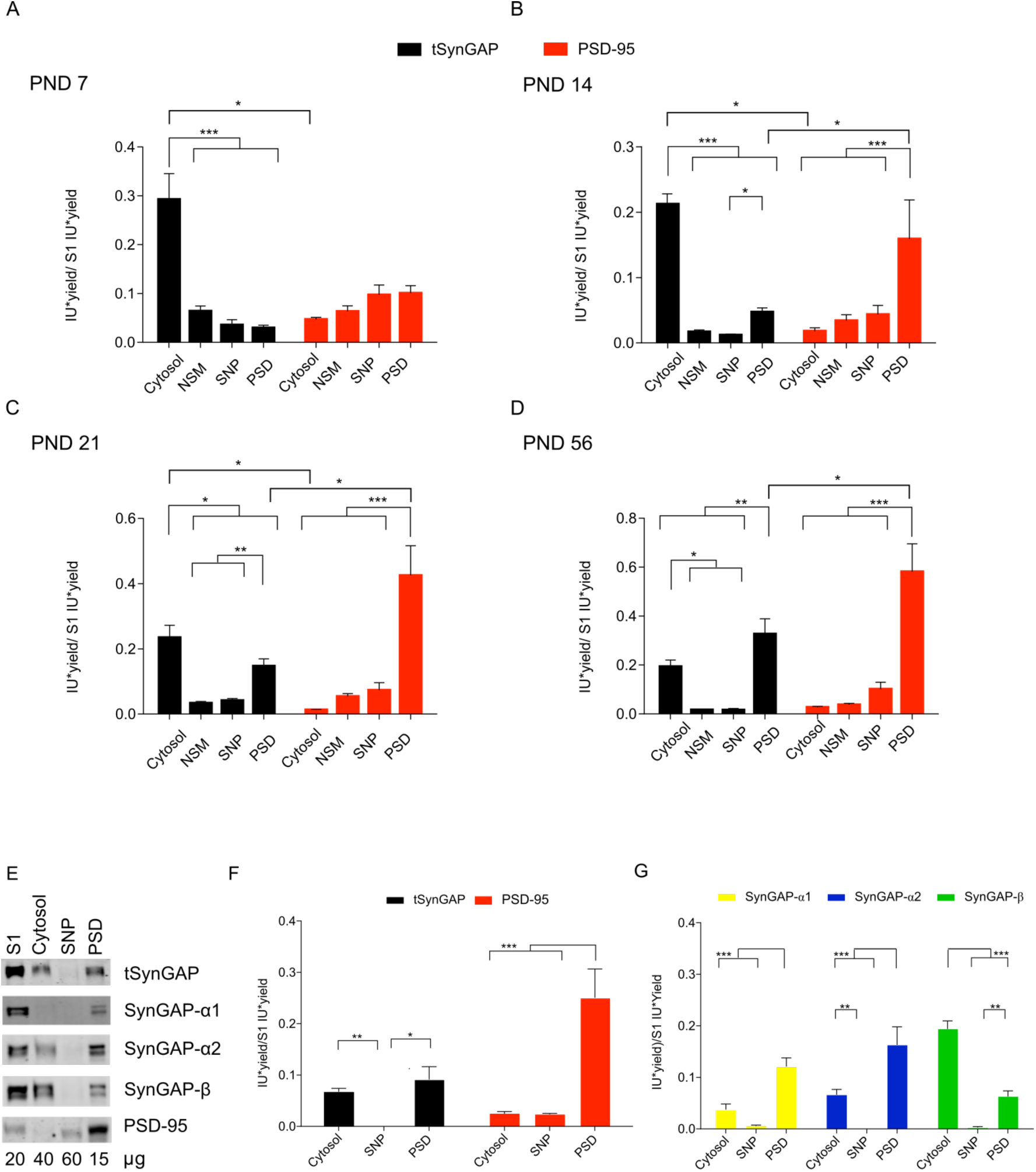
Subcellular distribution of tSynGAP and PSD-95 in cortex from four life stages and adult hippocampus. **A-D.** Bar plots representing the mean of normalized immunoblot intensity data (see Supplementary Figure 3B) from different subcellular fractions. Black bars present total SynGAP data and red bars PSD-95 data. The standard error of the mean (SEM) is also shown. Mean differences were analyzed by one-way ANOVA followed by Tukey’s post-hoc test *** p < 0.001, ** p < 0.01 and * p < 0.05. Subcellular fractions correspond with: cytosol; NSM, non-synaptic membranes; SNP, synaptic non-PSD and PSD, postsynaptic density. Life stages investigated are: PND7 (A), PND14 (B), PND 21 (C) and PND56 (D). **E.** Immunoblots presenting subcellular distribution of total SynGAP (tSynGAP), its isoforms and PSD-95 in adult (PND56) hippocampus. Subcellular fractions investigated: S1, total homogenate without the nuclear fraction; cytosol; SNP, synaptic non-PSD and PSD, postsynaptic density. **F.** Bar plot with the mean of normalized immunoblot intensity data of tSynGAP (black bars) and PSD-95 (red bars) in the subcellular fractions obtained from adult hippocampus. **G.** Bar plot with mean of normalized immunoblot intensity data of SynGAP isoforms containing α1, α2 and β C-terminal variants in the subcellular fractions obtained from adult hippocampus. The standard error of the mean (SEM) is also shown. Mean differences were analyzed by one-way ANOVA followed by Tukey’s post-hoc test *** p < 0.001, ** p < 0.01 and * p < 0.05.

### Differential subcellular distribution of SynGAP isoforms along postnatal development

We also investigated the subcellular distribution of SynGAP isoforms along postnatal development. As expected, in this study we also localize all isoforms at two main subcellular locations, the cytosol and the PSD. Their presence in non-synaptic membranes or in synaptic locations other than the PSD is almost negligible in all life stages investigated (Fig. 6 and 7). We found very low levels of α1 isoforms early in postnatal development (PND7 & 14), and as a consequence, their subcellular location could not be confidently established at these ages. At later ages, α1 isoforms presented a very restricted localization at the PSD (Fig. 7C,D). Alpha2- and β-containing isoforms were expressed at higher levels early in development and could thus be localized at specific subcellular locations from PND7 to 56. These isoforms presented and almost exclusive cytosolic location during the two first postnatal weeks (Fig. 7A,B), and a much-increased PSD localization between PND21 and PND56. Nevertheless, β isoforms were always found significantly more expressed in the cytosol than in the PSD, even at PND56. In contrast, α2 isoforms present the same expression level in cytosol and PSD at both PND21 and 56, although PSD levels are slightly higher. We also investigated adult (PND56) subcellular localization of SynGAP and its isoforms in hippocampus (Fig. 6E-G). Here we could not find a significantly different expression level of tSynGAP between cytosol and PSD, while PSD-95 presented a very restricted PSD localization. Similar to what we observed in PND56 cortex, α1 and α2 isoforms are enriched at the PSD, while β isoforms are very much enriched in the cytosolic fraction.

**Figure 7.**
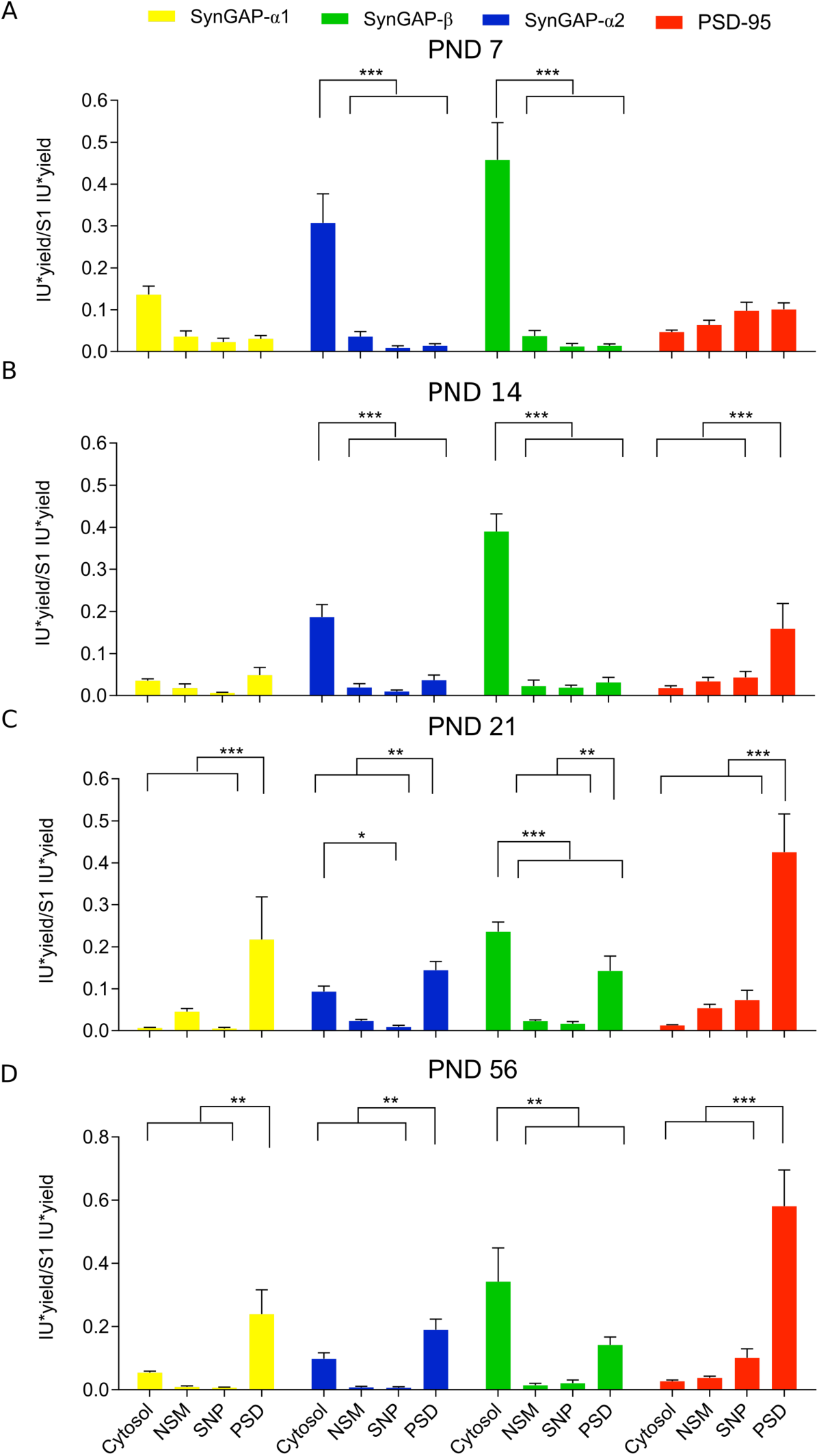
Subcellular distribution of SynGAP isoforms along three postnatal developmental stages and adulthood. **A-D.** Bar plots representing the mean of normalized immunoblot intensity data (see Supplementary Figure 3B) from different subcellular fractions for SynGAP isoforms presenting three different C-terminal variants and PSD-95. The standard error of the mean (SEM) is also shown. Mean differences were analyzed by one-way ANOVA followed by Tukey’s post-hoc test *** p < 0.001, ** p < 0.01 and * p < 0.05. Subcellular fractions correspond with: cytosol; NSM, non-synaptic membranes; SNP, synaptic non-PSD and PSD, postsynaptic density. Life stages investigated are: PND7 (A), PND14 (B), PND 21 (C) and PND56 (D).

## Discussion

It is established that mammals express multiple protein isoforms from the *Syngap1* gene, yet the complete set of SynGAP isoforms is still to be defined. Evidence from multiple sources, including transcriptomics data, suggest that human, mouse and rat SynGAP isoforms would at least have 5 N-termini (A1, A2, B, C and D) and 4 C-termini (α1, α2, β and γ). Nevertheless, C and D N-termini have not been reported in humans and NCBI adds four extra N-termini (A3, A4, E and F), and another C-terminus (α3), to the mouse set of variants. To the best of our knowledge, only the D N-terminus has been unambiguously identified at the protein level, which we report here for the first time. Originally described in rat at the RNA level (Li *et al*. 2001), the D variant was submitted into the NCBI as an artefactual sequence resulting from the fusion of transcripts from two different genes, as reported later (McMahon *et al*. 2012), and thus considered nonexistent. Yet, our proteomics experiments identify a peptide corresponding to the N-terminus of this variant. We have identified this peptide in cortical samples from all ages investigated (PND1-56). Importantly, the first methionine of this peptide is acetylated, a common post-translational modification of N-termini having a glutamate in the second position (Varland *et al*. 2015), as in the D variant. The recurrent identification of this acetylated N-terminal peptide provides strong evidence for the expression of this variant in mouse cortex. The fact that mass spectrometry-based methods have not been able to identify unique peptides from the other isoforms, beyond peptides common to A1 and A2 (McMahon *et al*. 2012), suggests that their N-termini could be proteolyzed.

Different SynGAP isoforms have been shown to have opposed functions in the control of synaptic strength (McMahon *et al*. 2012), to localize to different subcellular compartments (Li *et al*. 2001; Moon *et al*. 2008; Tomoda 2004) or to be able to bind to different proteins (Li *et al*. 2001; Kim *et al*. 1998), participating in different protein complexes and, by extension, molecular functions. SynGAP-β has even been described at the nucleus of cortical and hippocampal neurons (Moon *et al*. 2008), which would be in agreement with the nuclear localization signal that the Database of Nuclear Localization Signals (Bernhofer *et al*. 2017) identifies in the SynGAP core region (KKRKKD). This motif would be located towards the end of the C2 domain, as it occurs in the Doc2g protein (Fukuda *et al*. 2001). It is for this reason that it is essential to understand where and when these isoforms are expressed to have a correct understanding of SynGAP biology. Using three highly specific antibodies against α1, α2 and β C-termini we compared isoform expression levels in different brain regions along postnatal development and in adult, as well as their cortical and hippocampal subcellular distribution. We have also shown that isoforms presenting these three C-termini are also expressed in the cortex of two human individuals aged 19 and 67. As previously reported (Kim *et al*. 1998; Li *et al*. 2001; McMahon *et al*. 2012; Yang *et al*. 2013), the three antibodies identify two major bands in all conditions investigated, including in human samples. This indicates that each C-terminus will at least be expressed with two different N-termini. Importantly, we did not observe different expression patterns between the top and bottom bands, indicating that the C-termini strongly determine the differential expression observed between isoforms.

Developmental expression in each of the five tissues analyzed revealed important differences between isoforms. These were particularly clear in cortex, hippocampus and striatum. As in cerebellum and olfactory bulb the levels observed at PND4 remained unaltered through adulthood, for β isoforms, or just increased moderately for α1 and α2 isoforms. In cortex and hippocampus, the temporal expression of α1 isoforms is importantly delayed relative to that of α2 and β isoforms, as α1 isoforms reach their maximum at PND56, while α2 and β at PND21. Indeed, α1 isoforms present very low levels at PND11, during the critical period of *Syngap1* deficiency (Clement *et al*. 2012; Clement *et al*. 2013), but later experience a rapid increase in their expression. For instance, between PND11 and 21, α1 isoforms increase their abundance 5-fold in cortex. The developmental expression observed in striatum presents a bi-phasic pattern, which is unique to this brain region. Protein expression increases between PND4 and PND11, remains constant during the critical period, between PND11 and 21, and increases again between PND21 and PND56. This pattern in very obvious for α1 and α2 isoforms, while the second stage of increase, between PND21 and PND56, does not reach statistical significance for β isoforms. Taken together, these data support the idea of different functionalities of SynGAP isoforms.

As PSD-95 is known to interact with α1 isoforms and CaMKIIα with β ones, we also investigated if the developmental expression changes observed for these two pairs of proteins were correlated and, if so, in what tissues. We found a clear correlation for the developmental expression of PSD-95 and α1 isoforms in cortex, but also in hippocampus, striatum (p = 0.06) and olfactory bulb. The expression of β isoforms with CaMKIIα only correlated in cortex and hippocampus. Interestingly, the cortical decreases in β isoforms abundance occurring between PND21 and 56 was also observed for CaMKIIα. No correlation was found between α1 isoforms and CaMKIIα or even between α1 and β isoforms in any of the tissues investigated.

Expression comparison between tissues revealed that cerebellum and olfactory bulb presented lower expression levels for all isoforms and ages, with the exception of α2 and β isoforms in the olfactory bulb and at PND4. Isoform abundance comparison between cortex, hippocampus and striatum revealed that at PND11 and adulthood the highest expression was found in hippocampus. Yet, at PND21 clear differences where observed between isoforms. At this stage, α1 isoforms presented the highest expression levels at the cortex, α2 isoforms presented equivalent levels in cortex and hippocampus, both being higher than striatum, while hippocampus remained the tissue with the highest expression of β isoforms. We thus observed an increase in the cortical expression of α1 and α2 isoforms relative to their hippocampal levels, this being particularly striking for α1 isoforms. Importantly, the changes in isoform expression observed between PND14 and 21, take place during the *Syngap1* critical period (Clement *et al*. 2012), suggesting that they could have a role in disease. Remarkably, after this period, the expression pattern returns to that observed at PND11. For this reason, PND21 could be a key age for SynGAP neurobiology.

As SynGAP was first identified in the PSD, it has mainly been studied in the context of adult synaptic function. Yet, *Syngap1* expression starts early in embryogenesis (Porter *et al*. 2005), before the onset of synaptogenesis, indicating that SynGAP must have non-synaptic functions. For this reason, we decided to investigate the distribution of SynGAP into subcellular compartments and if this changed along postnatal development. We looked for SynGAP distribution in 4 major cellular compartments, i) cytosol, containing soluble proteins not bound to membranes, ii) non-synaptic membranes (NSM), iii) synapse excluding the PSD (SNP) and iv) the PSD, and we compared it to the distribution of the PSD marker PSD-95. Although we detected SynGAP and its isoforms in all subcellular fractions investigated, when normalizing their abundance by the amount of protein found in each fraction, we observed that SynGAP largely partitions between two locations, the PSD and the cytosol. Indicating that SynGAP has a cytosolic function, which is essentially uncharacterized, and that this is the predominant function early in postnatal development.

tSynGAP subcellular distribution was remarkably different from that of PSD-95, which at PND7 was weakly expressed, but from PND14 onwards was almost exclusively found at the PSD. Instead, tSynGAP presented a clear cytosolic localization at all ages investigated, even in adult cortex and hippocampus. Actually, during the first two postnatal weeks, tSynGAP was almost exclusively found at the cytosol. The fraction of tSynGAP localized to the PSD progressively increased along postnatal development but tSynGAP was only found enriched in the PSD in adult cortical samples. Even at PND21, when synaptogenesis is largely finished (Harris 1999), tSynGAP was still found more abundant in the cytosol. It is also interesting to note that within synapotosomes, SynGAP is almost exclusively found at the PSD, as we detected very low amounts of this protein at the SNP fraction, which contains all synaptic proteins that are not in the PSD. Different SynGAP isoforms presented a clearly distinctive subcellular localization pattern between the cytosol and the PSD. Being α1 and β isoforms the ones with the most opposed patterns. Alpha1 isoforms were always found highly restricted to the PSD, while β ones were always enriched in the cytosolic fraction and absent from the PSD until PND21. Isoforms with the α2 C-termini presented an intermediate behavior, enriched in the cytosol in PND7 and 14 and enriched at the PSD at PND21 and 56. This differential distribution of SynGAP isoforms was also observed in human hippocampal samples, indicating that the same localization pattern could be found in other brain regions.

Interestingly, at PND14, when PSDs are already formed in the mouse cortex (Swulius *et al*. 2010; Chandrasekaran *et al*. 2015), as indicated by the clear enrichment of PSD-95 in this fraction, α2 and β containing isoforms present very low PSD levels. However, one week later (PND21), coinciding with the rise of α1 isoforms expression and localization to the PSD, the presence of α2 and β isoforms in this location clearly increases. This finding suggests that, an increase in PSD abundance of α1 isoforms would result in increased PSD amounts of the other isoforms. As α1 isoforms are very much restricted to the PSD, their presence there could help stabilize the other isoforms.

Taking into consideration that the interaction between SynGAP-α1 and PSD-95 is not required for SynGAP localization to the PSD (Barnett *et al*. 2006; Vazquez *et al*. 2004; Rumbaugh *et al*. 2006) we propose a model in which all SynGAP isoforms would be able to locate to the PSD through a primary interaction with a yet unidentified protein. This interaction would occur via a sequence within the core region of SynGAP. Furthermore, α1 isoforms would present a secondary PSD anchoring point at their PDZ binding motif, which would result in their increased stability at the PSD. This increased stabilization would not go in detriment of the well-described dispersion of α1 and α2 isoforms from the PSD upon synaptic activation, as dispersed isoforms rapidly return to the PSD (Yang *et al*. 2011; Yang *et al*. 2013; Araki *et al*. 2015). Furthermore, the low abundance of α1 isoforms in the cytosol suggests that at any given time the proportion of α1 isoforms outside the PSD is relatively small. Additionally, this model also proposes that the presence of α1 isoforms at the PSD would help stabilize α2 and β isoforms in this location. The interaction between isoforms with different C-termini would occur through the coiled-coil domain, which has been described to promote SynGAP oligomerization (Zeng *et al*. 2016). Alpha1 and α2 present an identical coiled coil domain, yet β isoforms present a shorter coiled-coil domain, which might result in a less efficient oligomerization (Zeng *et al*. 2016). If this is the case, it might explain why β isoforms remain more abundant in the cytosol. Nevertheless, it can not be ruled out that between PND14 and PND21 the primary and yet uncknown interacting point for SynGAP at the PSD becomes much more abundant, driving the increase of α2 and β isoforms at this location, instead of its interaction with α1 isoforms.

In summary, we have identified clear developmental expression pattern differences between SynGAP isoforms, particularly during the critical period of *Syngap1* haploinsufficiency, which could have relevance to brain development and mental illness. Furthermore, we provide strong evidence showing that SynGAP, generally regarded as a protein exclusive to the PSD, is also found in the cytosol, where it is most abundant during postnatal brain development. Different isoforms present clearly distinctive subcellular distribution, being α1 isoforms highly restricted to the PSD, β ones mainly cytosolic and α2 isoforms presenting an intermediate behavior. We have observed that the presence of α2 and β isoforms in the PSD is developmentally regulated and coincides with the increased expression of α1 isoforms. Understanding the functional differences between these isoforms will be key to disentangle the multiple functions performed by the SynGAP protein.

## Acknowledgments

Financial support for this work was provided by: BFU2012-34398 and BFU2015-69717-P (MINECO), Career Integration Grant (ref. 304111), Ramón y Cajal Fellowship (RYC-2011-08391p), IEDI-2017-00822; AGAUR (SGR14-297 and 2017 SGR 1776) to AB; BES-2013-063720 (MINECO) to GG; MH096847 (NIH), MH108408 (NIH) and NS064079 (NIH) to GR and RO1 MH112151 (NIH) to RLH.

## Supplementary Figures and Tables

**Supplementary Table 1. Mouse, rat and human *Syngap1*/*SYNGAP1* protein isoforms identified by Ensembl, UniProt and NCBI Gene databases.**

**Supplementary Figure 1.**
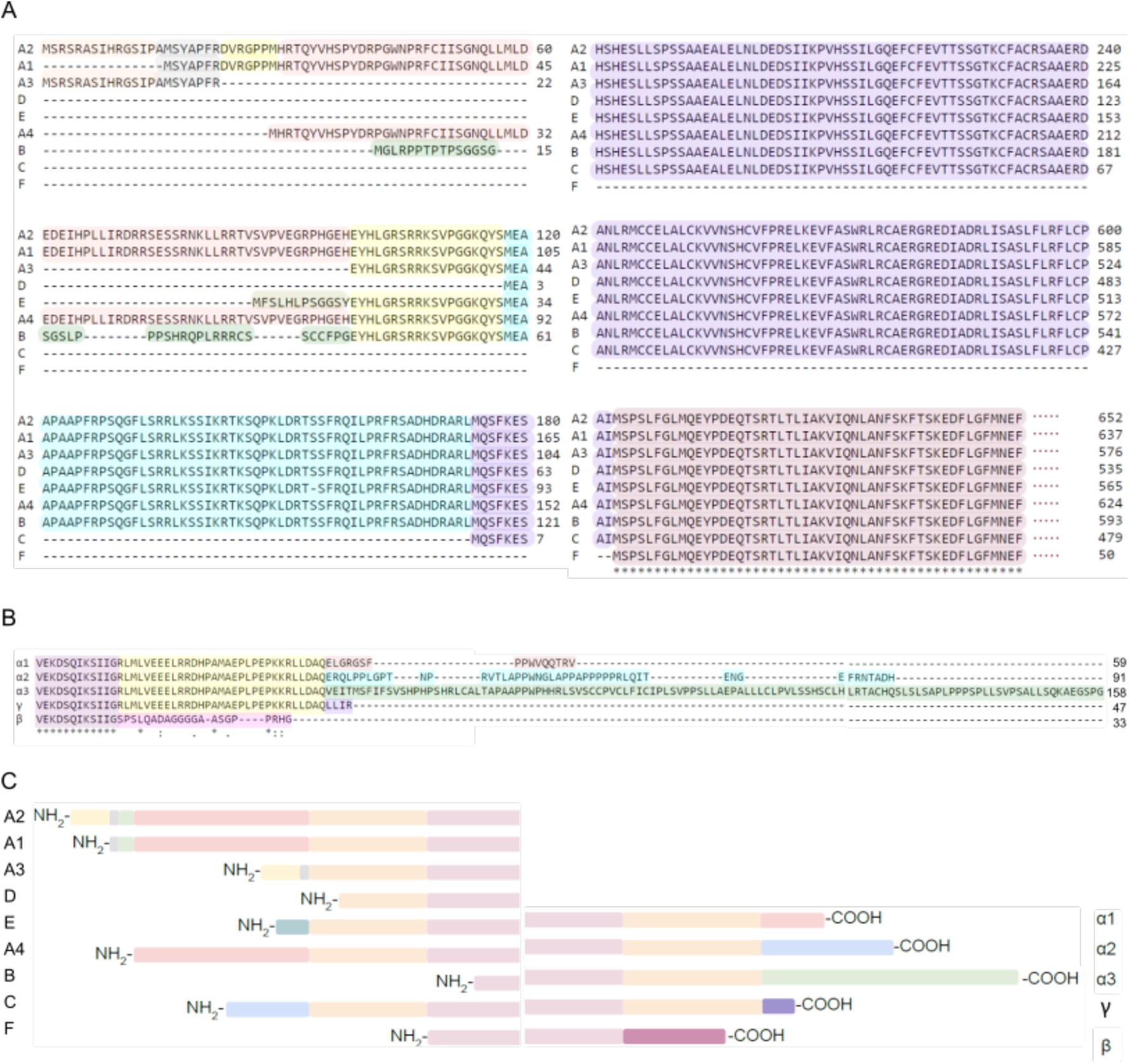
SynGAP N- and C-terminal variants. **A.** Protein sequences from mouse SynGAP protein N-terminal variants. **B.** Protein sequences from mouse SynGAP protein C-terminal variants. **C-D.** Schematic representation of SynGAP protein N- and C-terminal variants. Fragments with the same color correspond with the same protein sequence. The beginning and the end of the SynGAP core region, that is the sequence common to all isoforms, is also indicated.

**Supplementary Table 2. N- and C-terminal variants identified by mass spectrometry from mouse cortex.**

**Supplementary Figure 2.**
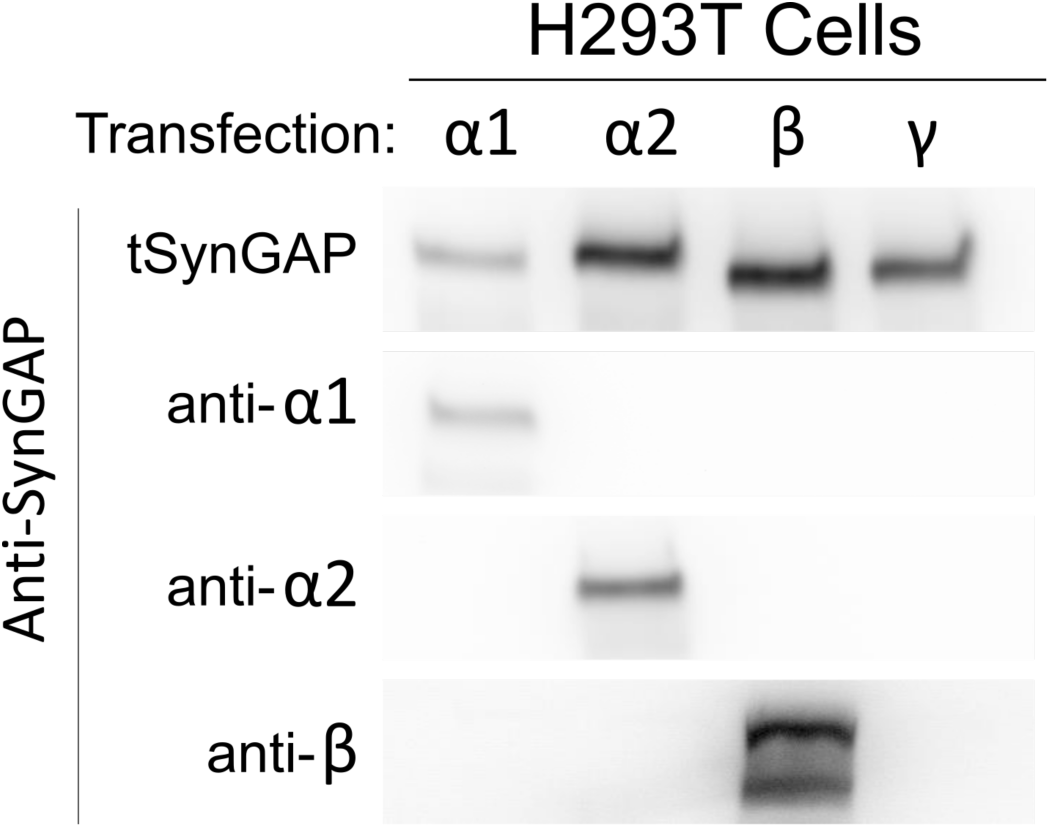
Specificity of anti-SynGAP-β antibody. Pools of HEK293T cells were transfected with GFP-tagged SynGAP cDNAs containing one of the four known C-terminal spliced sequences. Extracts from each cellular pool were immunoblotted with and antibody that was common to all SynGAP isoforms (pan-SynGAP) or antibodies raised against α1, α2, and β spliced variants.

**Supplementary Figure 3.**
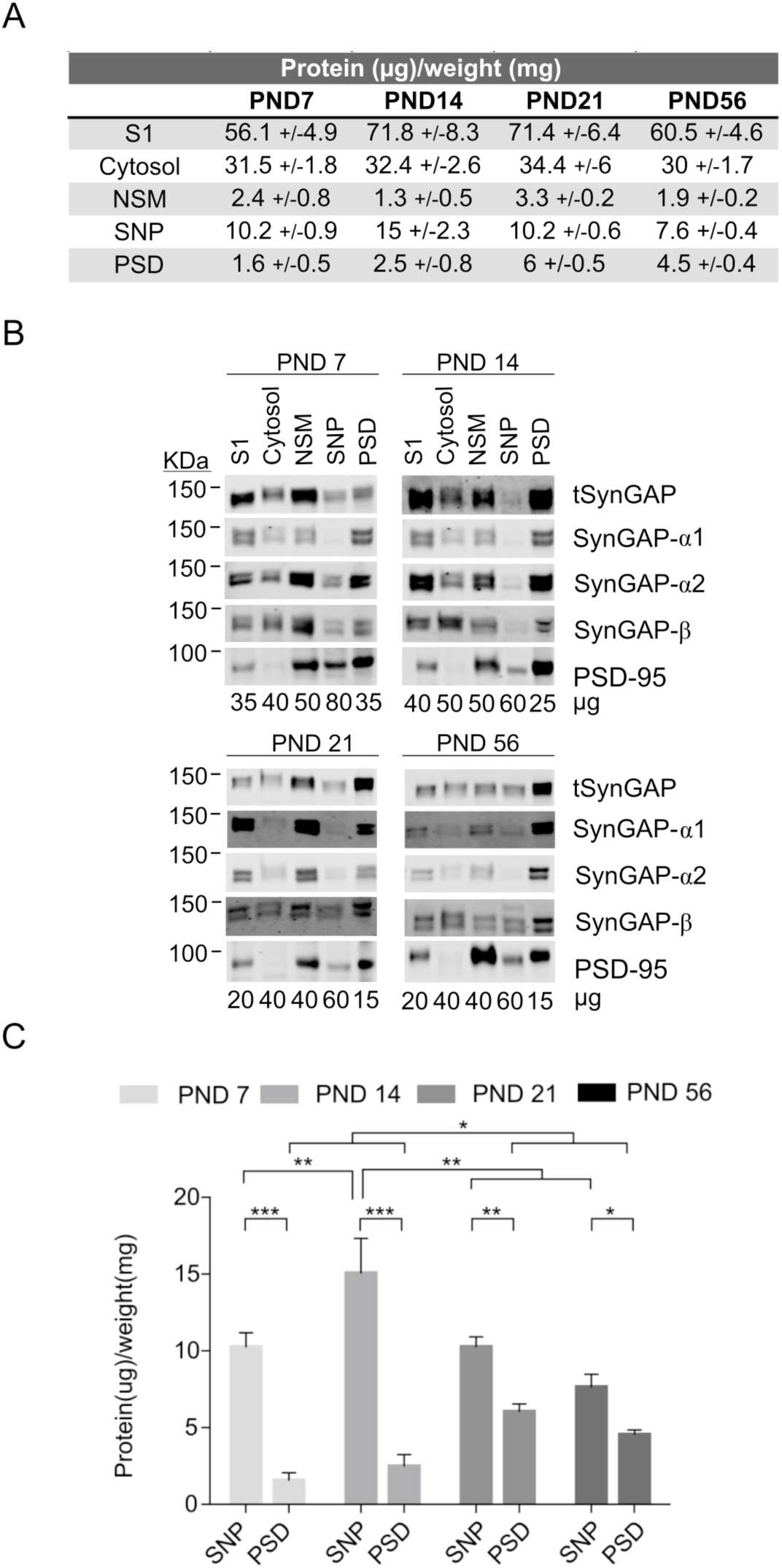
Subcellular fractionation of mouse cortex and immunoblot analysis of SynGAP and its isoforms expression in four life stages. **A.** Table presenting protein yields from all subcellular fractions generated. Protein yield is calculated as the ratio of total protein (in µg) in a given fraction by the weight (in mg) of the tissue used to obtain it. Subcellular fractions produced are: S1, total homogenate without the nuclear fraction; cytosol; NSM, non-synaptic membranes; SNP, synaptic not PSD and PSD, postsynaptic density. **B.** Representative immunoblots of total SynGAP (tSynGAP), its isoforms and PSD-95 in the subcellular fractions produced are shown for each life stage analyzed. Protein amounts (µg) used for each fraction in immunoblots is indicated below each lane. **C.** Bar plots represent protein yield for SNP and PSD fractions at each of the life stages investigated (PND7, 14, 21 and 56, N: 6 for all ages). The standard error of the mean (SEM) is also shown. Statistical test used one-way ANOVA followed by Tukey’s post-hoc test *** p < 0.001, ** p < 0.01 and * p < 0.05.

## Notes

#### Summary of Updates

The second version of this manuscript only includes minor changes in the text, with no changes in its results, figures or conclusions

## References

Aceti M., Creson T. K., Vaissiere T., Rojas C., Huang W.-C., Wang Y.-X., Petralia R. S., Page D. T., Miller C. A., Rumbaugh G. (2014) Author’s Accepted Manuscript. BPS 77, 805–815.

Araki Y., Zeng M., Zhang M., Huganir R. L. (2015) Rapid Dispersion of SynGAP from Synaptic Spines Triggers AMPA Receptor Insertion and Spine Enlargement during LTP. Neuron 85, 173–189.

Barnett M. W., Watson R. F., Vitalis T., Porter K., Komiyama N. H., Stoney P. N., Gillingwater T. H., Grant S. G. N., Kind P. C. (2006) Synaptic Ras GTPase activating protein regulates pattern formation in the trigeminal system of mice. J Neurosci 26, 1355–1365.

Bayés A., Collins M. O., Croning M. D. R., Van De Lagemaat L. N., Choudhary J. S., Grant S. G. N. (2012) Comparative study of human and mouse postsynaptic proteomes finds high compositional conservation and abundance differences for key synaptic proteins. PLoS ONE 7, e46683.

Bernhofer M., Goldberg T., Wolf S., Ahmed M., Zaugg J., Boden M., Rost B. (2017) NLSdb—major update for database of nuclear localization signals and nuclear export signals. Nucleic Acids Research 46, D503–D508.

Berryer M. H., Hamdan F. F., Klitten L. L., Møller R. S., Carmant L., Schwartzentruber J., Patry L., et al. (2013) Mutations in SYNGAP1Cause Intellectual Disability, Autism and a Specific form of Epilepsy by Inducing Haploinsufficiency. Human Mutation, n/a–n/a.

Carlin R. K., Grab D. J., Cohen R. S., Siekevitz P. (1980) Isolation and characterization of postsynaptic densities from various brain regions: enrichment of different types of postsynaptic densities. J Cell Biol 86, 831–845.

Carlisle H. J., Manzerra P., Marcora E., Kennedy M. B. (2008) SynGAP regulates steady-state and activity-dependent phosphorylation of cofilin. J Neurosci 28, 13673–13683.

Chandrasekaran S., Navlakha S., Audette N. J., McCreary D. D., Suhan J., Bar-Joseph Z., Barth A. L. (2015) Unbiased, High-Throughput Electron Microscopy Analysis of Experience-Dependent Synaptic Changes in the Neocortex. Journal of Neuroscience 35, 16450–16462.

Chen H. J., Rojas-Soto M., Oguni A., Kennedy M. B. (1998) A synaptic Ras-GTPase activating protein (p135 SynGAP) inhibited by CaM kinase II. Neuron 20, 895–904.

Clement J. P., Aceti M., Creson T. K., Ozkan E. D., Shi Y., Reish N. J., Almonte A. G., et al. (2012) Pathogenic SYNGAP1 Mutations Impair Cognitive Development by Disrupting Maturation of Dendritic Spine Synapses. Cell 151, 709–723.

Clement J. P., Ozkan E. D., Aceti M., Miller C. A., Rumbaugh G. (2013) SYNGAP1 links the maturation rate of excitatory synapses to the duration of critical-period synaptic plasticity. Journal of Neuroscience 33, 10447–10452.

Creson T. K., Rojas C., Hwaun E., Vaissiere T., Kilinc M., Jimenez-Gomez A., Holder J. L., et al. (2019) Re-expression of SynGAP protein in adulthood improves translatable measures of brain function and behavior. Elife 8.

Defelipe J., Alonso-Nanclares L., Arellano J. I. (2002) Microstructure of the neocortex: comparative aspects. J Neurocytol 31, 299–316.

Fukuda M., Saegusa C., Kanno E., Mikoshiba K. (2001) The C2A Domain of Double C2 Protein γ Contains a Functional Nuclear Localization Signal. Journal of Biological Chemistry 276, 24441–24444.

Hamdan F. F., Gauthier J., Spiegelman D., Noreau A., Yang Y., Pellerin S., Dobrzeniecka S., et al. (2009) Mutations in SYNGAP1 in autosomal nonsyndromic mental retardation. N. Engl. J. Med. 360, 599–605.

Harris K. M. (1999) Structure, development, and plasticity of dendritic spines. Current Opinion in Neurobiology 9, 343–348.

Jeyabalan N., Clement J. P. (2016) SYNGAP1: Mind the Gap. Front. Cell. Neurosci. 10, 32.

Kilinc M., Creson T., Rojas C., Aceti M., Ellegood J., Vaissiere T., Lerch J. P., Rumbaugh G. (2018) Species-conserved SYNGAP1 phenotypes associated with neurodevelopmental disorders. Mol Cell Neurosci 91, 140–150.

Kim J. H., Lee H.-K., Takamiya K., Huganir R. L. (2003) The role of synaptic GTPase-activating protein in neuronal development and synaptic plasticity. Journal of Neuroscience 23, 1119–1124.

Kim J. H., Liao D., Lau L. F., Huganir R. L. (1998) SynGAP: a synaptic RasGAP that associates with the PSD-95/SAP90 protein family. Neuron 20, 683–691.

Klooster ten J. P., Hordijk P. L. (2007) Targeting and localized signalling by small GTPases. Biol. Cell 99, 1–12.

Komiyama N. H., Watabe A. M., Carlisle H. J., Porter K., Charlesworth P., Monti J., Strathdee D. J., et al. (2002) SynGAP regulates ERK/MAPK signaling, synaptic plasticity, and learning in the complex with postsynaptic density 95 and NMDA receptor. J Neurosci 22, 9721–9732.

Krapivinsky G., Medina I., Krapivinsky L., Gapon S., Clapham D. E. (2004) SynGAP-MUPP1-CaMKII synaptic complexes regulate p38 MAP kinase activity and NMDA receptor-dependent synaptic AMPA receptor potentiation. Neuron 43, 563–574.

Li W., Okano A., Tian Q. B., Nakayama K., Furihata T., Nawa H., Suzuki T. (2001) Characterization of a Novel synGAP Isoform, synGAP-beta. Journal of Biological Chemistry 276, 21417–21424.

McMahon A. C., Barnett M. W., O’Leary T. S., Stoney P. N., Collins M. O., Papadia S., Choudhary J. S., et al. (2012) SynGAP isoforms exert opposing effects on synaptic strength. Nat Comms 3, 900.

Michaelson S. D., Ozkan E. D., Aceti M., Maity S., Llamosas N., Weldon M., Mizrachi E., et al. (2018) SYNGAP1 heterozygosity disrupts sensory processing by reducing touch-related activity within somatosensory cortex circuits. Nat. Neurosci. 21, 1–13.

Mignot C., Stülpnagel von C., Nava C., Ville D., Sanlaville D., Lesca G., Rastetter A., et al. (2016) Genetic and neurodevelopmental spectrum of SYNGAP1-associated intellectual disability and epilepsy. Journal of Medical Genetics 53, 511–522.

Moon I. S., Sakagami H., Nakayama J., Suzuki T. (2008) Differential distribution of synGAP alpha1 and synGAP beta isoforms in rat neurons. Brain Res. 1241, 62–75.

Ozkan E. D., Creson T. K., Kramár E. A., Rojas C., Seese R. R., Babyan A. H., Shi Y., et al. (2014) Reduced Cognition in Syngap1 Mutants Is Caused by Isolated Damage within Developing Forebrain Excitatory Neurons. Neuron 82, 1317–1333.

Parker M. J., Fryer A. E., Shears D. J., Lachlan K. L., McKee S. A., Magee A. C., Mohammed S., et al. (2015) De novo, heterozygous, loss-of-function mutations in SYNGAP1 cause a syndromic form of intellectual disability. Am J Med Genet A 167A, 2231–2237.

Pena V., Hothorn M., Eberth A., Kaschau N., Parret A., Gremer L., Bonneau F., Ahmadian M. R., Scheffzek K. (2008) The C2 domain of SynGAP is essential for stimulation of the Rap GTPase reaction. EMBO Rep. 9, 350–355.

Porter K., Komiyama N. H., Vitalis T., Kind P. C., Grant S. G. N. (2005) Differential expression of two NMDA receptor interacting proteins, PSD-95 and SynGAP during mouse development. Eur J Neurosci 21, 351–362.

Qin Y., Zhu Y., Baumgart J. P., Stornetta R. L., Seidenman K., Mack V., Van Aelst L., Zhu J. J. (2005) State-dependent Ras signaling and AMPA receptor trafficking. Genes & Development 19, 2000–2015.

Rumbaugh G., Adams J. P., Kim J. H., Huganir R. L. (2006) SynGAP regulates synaptic strength and mitogen-activated protein kinases in cultured neurons. Proc Natl Acad Sci USA 103, 4344–4351.

Spijker S. (2011) Dissection of Rodent Brain Regions, in Neuromethods, Vol. 57, pp.13–26. Humana Press, Totowa, NJ.

Swulius M. T., Kubota Y., Forest A., Waxham M. N. (2010) Structure and composition of the postsynaptic density during development. J Comp Neurol 518, 4243–4260.

Tomoda T. (2004) Role of Unc51.1 and its binding partners in CNS axon outgrowth. Genes & Development 18, 541–558.

Varland S., Osberg C., Arnesen T. (2015) N-terminal modifications of cellular proteins: The enzymes involved, their substrate specificities and biological effects. Proteomics 15, 2385–2401.

Vazquez L. E., Chen H. J., Sokolova I., Knuesel I., Kennedy M. B. (2004) SynGAP regulates spine formation. Journal of Neuroscience 24, 8862–8872.

Vlaskamp D. R. M., Shaw B. J., Burgess R., Mei D., Montomoli M., Xie H., Myers C. T., et al. (2019) SYNGAP1 encephalopathy: A distinctive generalized developmental and epileptic encephalopathy. Neurology 92, e96–e107.

Walkup W. G., Sweredoski M. J., Graham R. L., Hess S., Kennedy M. B. (2018) Phosphorylation of synaptic GTPase-activating protein (synGAP) by polo-like kinase (Plk2) alters the ratio of its GAP activity toward HRas, Rap1 and Rap2 GTPases. Biochem. Biophys. Res. Commun. 503, 1599–1604.

Walkup W. G., Washburn L. R., Sweredoski M. J., Carlisle H. J., Graham R. L., Hess S., Kennedy M. B. (2014) Phosphorylation of Synaptic GTPase Activating Protein (synGAP) by Ca2+/Calmodulin-dependent Protein Kinase II (CaMKII) and Cyclin-dependent Kinase 5 (CDK5) Alters the Ratio of its GAP Activity Toward Ras and Rap GTPases. J. Biol. Chem.

Yang Y., Tao-Cheng J. H., Reese T. S., Dosemeci A. (2011) SynGAP Moves out of the core of the postsynaptic density upon depolarization. Neuroscience 192, 132–139.

Yang Y., Tao-Cheng J.-H., Bayer K. U., Reese T. S., Dosemeci A. (2013) Camkii-mediated phosphorylation regulates distributions of Syngap-α1 and -α2 at the postsynaptic density. PLoS ONE 8, e71795.

Zeng M., Shang Y., Araki Y., Guo T., Huganir R. L., Zhang M. (2016) Phase Transition in Postsynaptic Densities Underlies Formation of Synaptic Complexes and Synaptic Plasticity. Cell 166, 1163–1175.e12.

Zhu J. J., Qin Y., Zhao M., Van Aelst L., Malinow R. (2002) Ras and Rap control AMPA receptor trafficking during synaptic plasticity. Cell 110, 443–455.

