## Supplementary Table 2 for "SynGAP Splice Variants Display Heterogeneous Spatio-Temporal Expression And Subcellular Distribution In The Developing Mammalian Brain"

| Age | SynGAP variant | Unique peptide sequence | Precursor | Area |
|---|---|---|---|---|
| PND 0/1 | SynGAP-D | mEAAPAAPFRPSQGFLSR | 995.9938++ | 1802994 |
| | SynGAP-D | mEAAPAAPFRPSQGFLSR | 664.3316+++ | 1885129 |
| | SynGAP-α2 | VTLAPPWNGLAPPAPPPPPR | 1023.0702++ | 3585347 |
| | SynGAP-α2 | VTLAPPWNGLAPPAPPPPPR | 682.3825+++ | 8029659 |
| | SynGAP-α2 | LQITENGEFR | 603.8093++ | 25052336 |
| | SynGAP-α2 | QLPPLGPTNPR | 595.3380++ | 273971392 |
| | SynGAP-β | SIIGSPSLQADAGGGGAASGPPR | 1012.0138++ | 42673360 |
| | SynGAP-β | SIIGSPSLQADAGGGGAASGPPR | 675.0116+++ | 16715653 |
| PND 11 | SynGAP-D | mEAAPAAPFRPSQGFLSR | 995.9938++ | 9702698 |
| | SynGAP-D | mEAAPAAPFRPSQGFLSR | 664.3316+++ | 6188801 |
| | SynGAP-α2 | QLPPLGPTNPR | 595.3380++ | 138484432 |
| | SynGAP-α2 | VTLAPPWNGLAPPAPPPPPR | 1023.0702++ | 15871170 |
| | SynGAP-α2 | VTLAPPWNGLAPPAPPPPPR | 682.3825+++ | 28712010 |
| | SynGAP-α2 | LQITENGEFR | 603.8093++ | 26766820 |
| | SynGAP-β | SIIGSPSLQADAGGGGAASGPPR | 1012.0138++ | 131803552 |
| | SynGAP-β | SIIGSPSLQADAGGGGAASGPPR | 675.0116+++ | 52839004 |
| PND 21 | SynGAP-D | mEAAPAAPFRPSQGFLSR | 995.9938++ | 16732052 |
| | SynGAP-D | mEAAPAAPFRPSQGFLSR | 664.3316+++ | 11927298 |
| | SynGAP-α2 | LQITENGEFR | 603.8093++ | 36717168 |
| | SynGAP-α2 | VTLAPPWNGLAPPAPPPPPR | 1023.0702++ | 6693875 |
| | SynGAP-α2 | VTLAPPWNGLAPPAPPPPPR | 682.3825+++ | 12733106 |
| | SynGAP-α2 | QLPPLGPTNPR | 595.3380++ | 588498688 |
| | SynGAP-β | SIIGSPSLQADAGGGGAASGPPR | 1012.0138++ | 136940752 |
| | SynGAP-β | SIIGSPSLQADAGGGGAASGPPR | 675.0116+++ | 45366896 |
| PND 56 | SynGAP-D | mEAAPAAPFRPSQGFLSR | 995.9938++ | 64078996 |
| | SynGAP-D | mEAAPAAPFRPSQGFLSR | 664.3316+++ | 50273976 |
| | SynGAP-α1 | GSFPPWVQQTR | 651.8331++ | 11963907 |
| | SynGAP-α2 | VTLAPPWNGLAPPAPPPPPR | 1023.0702++ | 9078114 |
| | SynGAP-α2 | VTLAPPWNGLAPPAPPPPPR | 682.3825+++ | 15689616 |
| | SynGAP-α2 | QLPPLGPTNPR | 595.3380++ | 1868298624 |
| | SynGAP-β | SIIGSPSLQADAGGGGAASGPPR | 1012.0138++ | 64078996 |
| | SynGAP-β | SIIGSPSLQADAGGGGAASGPPR | 675.0116+++ | 50273976 |

α
